# Transcriptome analysis of *Drosophila suzukii* reveals molecular mechanisms conferring pyrethroid and spinosad resistance

**DOI:** 10.1101/2024.06.17.599459

**Authors:** Christine A. Tabuloc, Curtis R. Carlson, Fatemeh Ganjisaffar, Cindy C. Truong, Ching-Hsuan Chen, Kyle M. Lewald, Sergio Hidalgo, Nicole L. Nicola, Cera E. Jones, Ashfaq A. Sial, Frank G. Zalom, Joanna C. Chiu

## Abstract

*Drosophila suzukii* lay eggs in soft-skinned, ripening fruits, making this insect a serious threat to berry production. Since its 2008 introduction into North America, growers have used insecticides, such as pyrethroids and spinosads, as the primary approach for *D. suzukii* management, resulting in detections of insecticide resistance in this pest. This study sought to identify the molecular mechanisms conferring insecticide resistance in these populations. We sequenced the transcriptomes of two pyrethroid- and two spinosad-resistant isofemale lines. In both pyrethroid-resistant lines and one spinosad-resistant line, we identified overexpression of metabolic genes that are implicated in resistance in other insect pests. In the other spinosad-resistant line, we observed an overexpression of cuticular genes that have been linked to resistance. Our findings enabled the development of molecular diagnostics that we used to confirm persistence of insecticide resistance in California, U.S.A. To validate these findings, we leveraged *D. melanogaster* mutants with reduced expression of metabolic or cuticular genes that were found to be upregulated in resistant *D. suzukii* to demonstrate that these genes are involved in promoting resistance. This study is the first to characterize the molecular mechanisms of insecticide resistance in *D. suzukii* and provides insights into how current management practices can be optimized.

## Introduction

*Drosophila suzukii* (Matsumura), also known as spotted wing *Drosophila*, is an invasive, agricultural pest native to Southeast Asia. It was first detected in the continental United States in 2008 ^1, 2^. Since then, *D. suzukii* has been detected across North America ^3, 4^, Europe ^5, 6^, South America ^7, 8^, and Africa^9^. Female *D. suzukii* flies have a serrated ovipositor that enables them to lay eggs in soft-skinned, ripening fruit such as strawberries and caneberries ^4,10^. The ability of *D. suzukii* to infest ripening fruit as opposed to over-ripened fruit, like other *Drosophila* species, poses a unique threat to both fresh and processed berry production. In fact, in the first year after its detection, *D. suzukii* caused considerable damage to raspberries, blackberries, blueberries, and cherries leading Bolda et al. (2010) to estimate the annual potential economic impact of this pest in the Pacific Coast region to be US$421.5 million.

Current *D. suzukii* management strategies consist of intensive spray programs ^11, 12^ of several insecticides including pyrethroids and spinosyns ^13–15^. Pyrethroid insecticides target the insect nervous system by binding to the active site of a voltage-gated sodium channel (VGSC), encoded by the gene *paralytic* (*para*), causing overstimulation of the nervous system that results in excessive twitching and eventually death ^16^. Spinosyn insecticides, on the other hand, target the insect nervous system by binding to an allosteric site on the alpha 6 subunit of the nicotinic acetylcholine receptor (nAChRα6), resulting in overexcitation of motor neurons, paralysis, and eventually death ^17, 18^. Due to repeated insecticide exposure, its short generation time, and high fecundity, *D. suzukii* has great potential for developing insecticide resistance ^19^. To date, there are three reports in California, U.S.A. that have documented spinosad- ^20, 21^ and pyrethroid-resistance ^22^ in *D. suzukii*, but the exact mechanisms driving these adaptations have yet to be described.

Thus far, common molecular mechanisms conferring insecticide resistance characterized in other insect pest species include penetration resistance, metabolic resistance, and target-site resistance. Penetration resistance occurs when there is an overexpression of cuticular genes such as *cuticle protein 30 (cpr30)*, resulting in a less penetrable insect integument and therefore reduced dermal entry of the insecticide ^23^. Metabolic resistance is defined by an upregulation of metabolic enzymes that detoxify the insecticide prior to its binding to the target protein ^24^. The classes of metabolic genes implicated in insecticide resistance include *cytochrome P450s* (*cyp*) ^25^, *glutathione S-transferases* (*gst*) ^26^, *esterases* ^27^, and *heat shock proteins* (*hsp*) ^28^. Finally, target-site resistance occurs when a mutation within the target protein prevents the docking of the insecticide to its binding site ^29, 30^. It is currently unknown whether one or more of these mechanisms underlie resistance observed in *D. suzukii* populations in California. Therefore, there is a need for identifying the possible molecular mechanisms conferring insecticide resistance in this pest as this will enable the development of molecular diagnostics to monitor insecticide resistance development in the field as well as provide insights into how growers can adjust their *D. suzukii* management programs.

Here, we generated isofemale lines from field-collected *D. suzukii* populations determined to be resistant to either pyrethroids (specifically zeta-cypermethrin) or spinosad. We then sequenced the transcriptomes of two zeta-cypermethrin-resistant and two spinosad-resistant isofemale lines and compared them to susceptible isofemale lines developed from the same field collections to investigate the molecular mechanisms driving insecticide resistance in these flies. We observed that *D. suzukii* flies resistant to zeta-cypermethrin exhibited elevated expression of genes encoding metabolic enzymes, indicative of metabolic resistance, as well as decreased expression of *para*. In one spinosad-resistant line, we observed evidence of metabolic resistance. However, we observed increased expression of cuticular genes in a second spinosad-resistant line, suggesting that penetration resistance is at play. Leveraging our results, we developed quantitative polymerase chain reaction (qPCR)-based assays to diagnose overexpression of specific target genes, which could reflect metabolic or penetration resistance. Our molecular analysis of two recently field-collected *D. suzukii* populations corroborated with insecticide bioassays to show that metabolic resistance is persisting in berry-growing regions in California. Finally, taking advantage of the genetic tools available in a closely related species *D. melanogaster,* we showed that *cyp* and *cpr* overexpression indeed promotes resistance to insecticides. Taken together, our findings showed that molecular analysis of these gene targets can be leveraged in efficient resistance development monitoring programs prior to conducting conventional insecticide bioassays, which are much more laborious. Our study also provides insights into how growers can adjust *D. suzukii* management protocols to counter insecticide resistance development.

## Results

### Development and identification of insecticide-resistant *Drosophila suzukii* isofemale lines

We first generated multiple isofemale lines derived from field-collected resistant *D. suzukii* populations in California, such that any observed gene expression differences between susceptible and resistant isofemale lines can be attributed to causal genetic determinants with higher confidence. This is more appropriate than comparing resistant and susceptible flies collected from geographically separated locations that may be more genetically dissimilar. Isofemale lines were developed from *D. suzukii* populations collected from either a strawberry or caneberry field in 2019 where an insecticide control failure was reported. The population collected from the strawberry field exhibited resistance to zeta-cypermethrin (herein referred to as “S” for strawberry) while the population collected from the caneberry field exhibited resistance to spinosad (herein referred to as “C” for caneberry). To identify susceptible and resistant isofemale lines developed from each of the two populations, discriminating dose bioassays were first performed on these lines together with control isofemale lines developed from a susceptible field population collected from an untreated orchard in California (herein referred to as Wolfskill)^21^ (**Fig. 1a-b; Suppl. Table 1-2**). Lines S1, S3, and S4 exhibited decreased mortality compared to Wolfskill when treated with zeta-cypermethrin (n=8) (**Fig. 1a; Suppl. Table 1**), while lines C3, C4, and C6 exhibited lower mortality when treated with spinosad (**Fig. 1b; Suppl. Table 2**). We opted to use two lines with the lowest rates of mortality per population for further analyses.

**Fig. 1:**
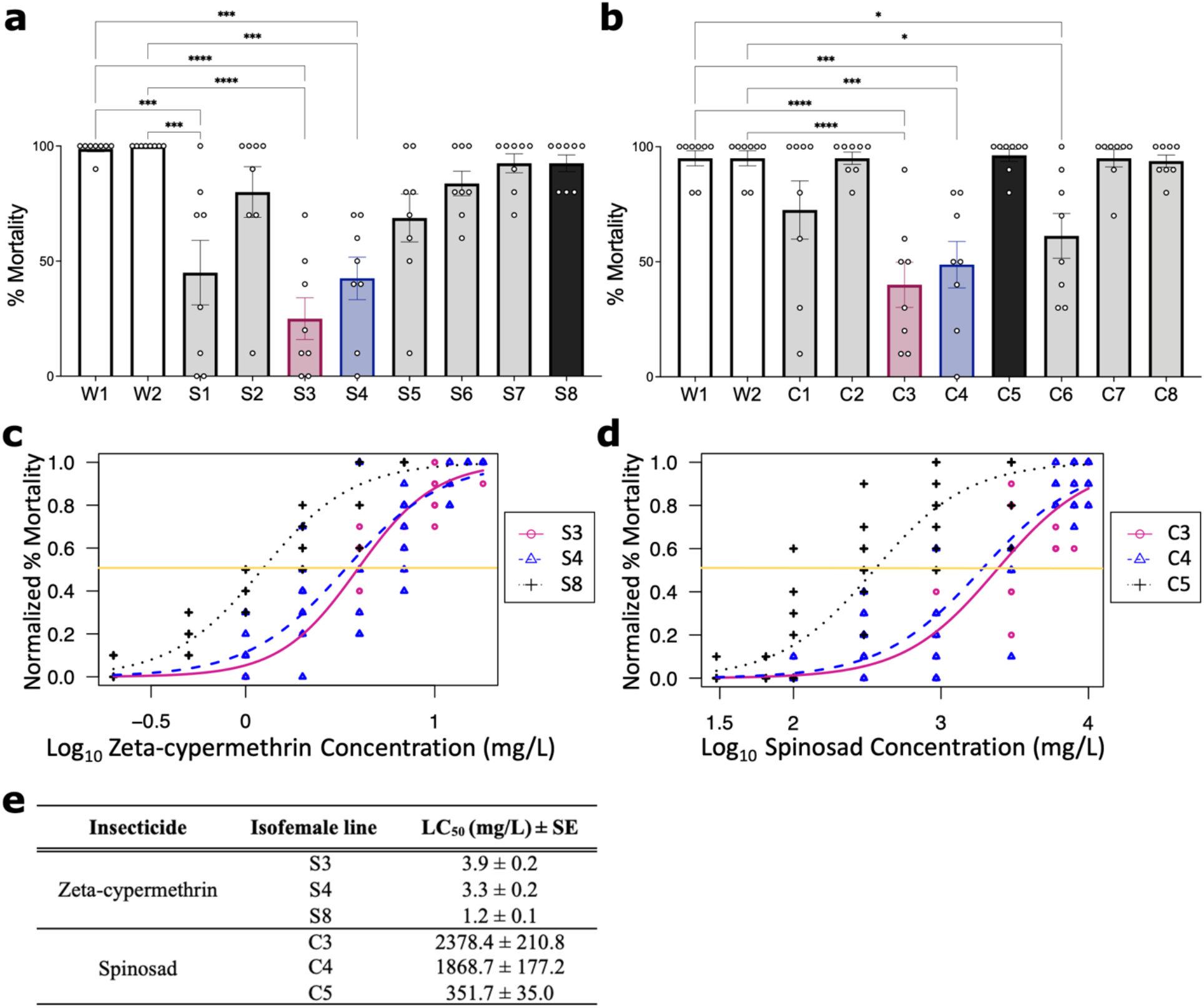
Identification of insecticide-resistant isofemale *D. suzukii* lines. (**a**-**b**) Bioassays to identify isofemale lines resistant to (**a**) zeta-cypermethrin (Mustang® Maxx) or (**b**) spinosad (Entrust® SC). Eight isofemale lines (indicated as S# and C#), developed from two separate field-resistant populations collected in California, USA, were tested. Two isofemale lines developed from a population collected from an untreated orchard in California, USA (Wolfskill, W#) served as the susceptible control (white). Each point represents a biological replicate of 5 males and 5 females (n=8), and error bars indicate ± SEM. Resistant lines used for subsequent experiments are indicated in maroon and blue while the susceptible line is in black. Asterisks denote significant p-values as determined by one-way ANOVA followed by Tukey’s multiple comparison test: * p<0.05, *** p<0.001, and **** p<0.0001. Non-significant comparisons are omitted. (**c**) Dose-response relationship between zeta-cypermethrin-resistant isofemale lines (S3: maroon circle and solid line; S4: blue triangle and dashed line) vs a susceptible sibling line (S8: black cross and dotted line) (n=8 biological replicates of 5 males and 5 females). The lethal concentration required to kill 50% of the population (LC_50_) is indicated by the yellow line. Each point represents a biological replicate. (**d**) Dose-response relationship between spinosad-resistant isofemale lines (C3: maroon circle and solid line; C4: blue triangle and dashed line) vs a susceptible sibling line (C5: black cross and dotted line) (n=8). (**e**) LC_50_ values of *D. suzukii* isofemale lines for zeta-cypermethrin and spinosad.

We next determined the effective concentration required to kill half of the tested population (LC_50_) of four resistant lines, two from each population (**Fig. 1c-1d**). The LC_50_ of the zeta-cypermethrin-resistant lines (S3, S4) were approximately three times greater than the susceptible line S8 derived from the same population (**Fig. 1c** and **e; Suppl. Table 3**). The LC_50_ of the spinosad-resistant lines (C3, C4) were at least five times higher than the susceptible line C5 derived from the same population (**Fig. 1d** and **e; Suppl. Table 3**). Therefore, we concluded that lines S3 and S4 are resistant to zeta-cypermethrin, and lines C3 and C4 are resistant to spinosad.

### Overexpression of metabolic genes suggests metabolic resistance in zeta-cypermethrin-resistant ***Drosophila suzukii***

We performed short-read RNA sequencing (RNA-Seq) to identify the molecular mechanisms underlying zeta-cypermethrin resistance in *D. suzukii*. We sequenced two zeta-cypermethrin-resistant lines (S3 and S4) and two susceptible lines derived from the same population as controls (S7 and S8). Pearson’s correlation confirmed that the biological replicates are highly correlated with one another (**Suppl. Fig. 1**).

To determine whether gene expression changes underlie insecticide resistance, we identified differentially expressed genes (DEGs) between the resistant and susceptible lines. We observed a total of 2,120 downregulated genes, 1,708 upregulated genes, and 8,723 non-differentially expressed genes between line S3 and the susceptible lines (**Fig. 2a; Suppl. Table 4**). For line S4, we identified 3,686 downregulated genes, 4,240 upregulated genes, and 6,323 non-differentially expressed genes (**Fig. 2b; Suppl. Table 5**). Amongst the upregulated genes are those encoding classes of metabolic enzymes known to confer insecticide resistance in other insect species. For instance, we observed that at least one of the two resistant lines exhibited a significant increase in the expression of *cytochrome P450* (*cyp*) *6a20* and *cyp4d14*, the carboxylesterase *cricklet*, *heat shock proteins* (*hsp*) *60B* and *hsp70Aa*, and *glutathione-s-transferase* (*gst*) *E3* (**Fig. 2c; Suppl. Table 6)**. Our results suggest that zeta-cypermethrin resistance in the original field-collected population may be attributed to metabolic resistance. Furthermore, many of the genes downregulated in the resistant lines are genes related to cellular signaling, such as nicotinic acetylcholine receptors, acetylcholine transporters, and voltage-gated sodium channel (VGSC) subunits; notably, the VGSC *paralytic* (*para*), the gene encoding the target protein of zeta-cypermethrin, is among them (**Fig. 2a-b, insert)**.

**Fig. 2:**
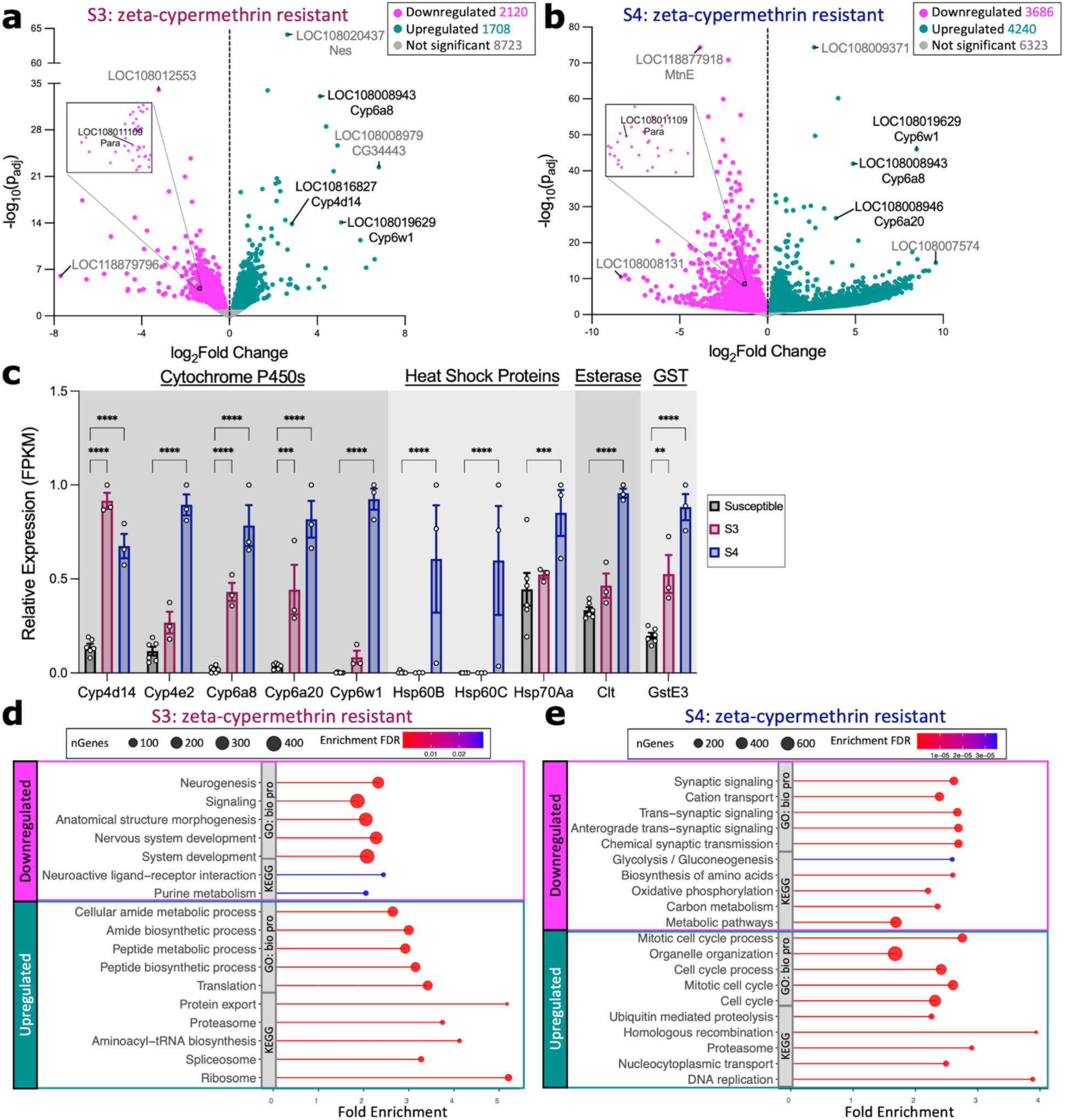
Zeta-cypermethrin-resistant lines exhibit increased expression of genes involved in metabolic resistance. (**a**) Volcano plot of genes displaying fold change gene expression differences between the resistant S3 line vs 2 susceptible lines (S7 and S8). Genes upregulated in the resistant populations (green) are to the right of the dotted line while downregulated genes (pink) lie to the left of the dotted line. Genes that exhibit no significant difference (NS) in expression between the two populations are in grey. Highly significant differences are higher up on the y-axis (where p_adj_ is the Benjamin-Hochberg adjusted p-value). Labeled points signify genes satisfying at least one of the following criteria: (1) have the largest fold change difference between the two groups, (2) have the lowest p_adj_, and (3) are genes known to be involved in insecticide resistance. Labels contain the *D. suzukii* gene symbol (LOC#########) and the corresponding *D. melanogaster* gene symbol homolog. Genes known to be involved in conferring resistance are labeled in black while genes that are not known to be involved in resistance are grey. The black box denotes the region containing *para*, zoomed-in in the insert. (**b**) Volcano plot of genes displaying fold change gene expression differences between the resistant S4 line vs 2 susceptible lines (S7 and S8). (**c**) Relative expression (FPKM) of *cytochrome P450* genes (*Cyp*), *heat shock proteins* (*Hsp*), the carboxylesterase *Cricklet* (*Clt*), and *glutathione-s-transferase E3* (*GstE3*) in the susceptible (S7 and S8: black) vs resistant (S3: maroon; S4: blue) groups extracted from the RNA sequencing data. Each point denotes a biological replicate (n=3 replicates of 8-10 females per line). Asterisks denote significant p-values as determined by 2-way ANOVA: ** p<0.01, ***p<0.001, and **** p<0.0001. (**d**-**e**) Top 5 enrichment pathways within the Kyoto Encyclopedia of Genes and Genomes (KEGG) and Gene Ontology (GO) Biological Processes (bio proc) categories for genes up- or down-regulated in line (**d**) S3 or (**e**) S4. The x-axis is Fold Enrichment, which is the percentage of differentially expressed genes that belong to each pathway. Point size represents the number of genes (nGenes) within the category while color denotes the false discovery rate (FDR) correction of enrichment p-values.

Functional enrichment analyses were performed to identify in which pathways these DEGs are involved in (**Fig. 2d-e; Suppl. Tables 7-8**). Downregulated genes in line S3 were enriched in several pathways involved in neuronal signaling while upregulated genes were enriched in pathways involved in RNA processing, protein expression, and metabolism (**Fig. 2d; Suppl. Table 7**). For line S4, downregulated genes were enriched in pathways involving neuronal signaling and metabolism while the upregulated genes are enriched in pathways involved in protein degradation and the cell cycle (**Fig. 2e; Suppl. Table 8**).

Next, we performed Weighted Gene Co-expression Network Analysis (WGCNA), an unsupervised analysis pipeline that clusters genes into modules based on their expression profile across samples ^31^, to identify potential novel gene clusters highly correlated with resistance (**Suppl. Fig. 2-3; Suppl. Table 9-10**). For line S3, genes were clustered into 35 different colored modules with turquoise being most correlated with resistance (R^2^ = 0.89) (**Suppl. Fig. 2a; Suppl. Table 9**). Of the 2,908 genes in turquoise, we identified several metabolic genes within the class of *cyp*s, *hsp*s, GSTs, and esterases (**Suppl. Fig. 2b**). Functional analysis revealed that the genes in turquoise are enriched in metabolic, RNA processing, and protein expression pathways (**Suppl. Fig. 2c; Suppl. Table 10)**. For line S4, genes were clustered into 27 modules, with turquoise being most correlated with resistance (R^2^ = 0.92) (**Suppl. Fig. 3a**). Of the 4,449 genes within turquoise, several are metabolic genes known to confer insecticide resistance in other species (**Suppl. Fig. 3b; Suppl. Table 11**). Genes in turquoise are enriched in RNA processing, cell cycle, cell differentiation, and protein and gene expression pathways (**Suppl. Fig. 3c; Suppl. Table 12**). Taken together, our results suggest that an upregulation of metabolic gene expression most likely confer zeta-cypermethrin resistance observed in *D. suzukii* in California.

### Overexpression of cuticular and metabolic genes suggests penetration and metabolic resistance in spinosad-resistant *Drosophila suzukii*

We sequenced the transcriptomes of two spinosad-resistant lines (C3, C4) and two susceptible lines (C2, C5) derived from the same population to determine the molecular mechanisms conferring spinosad-resistance. Pearson’s correlation coefficients revealed a strong correlation between biological replicates, confirming consistency between the replicates (**Suppl. Fig. 4**).

Next, we assessed gene expression differences between each resistant line vs both susceptible lines. For line C3, we observed 852 DEGs, with 492 downregulated genes, 360 upregulated genes, and 11,756 non-differentially expressed genes (**Fig. 3a; Suppl. Table 13**). In line C4, there were 4,233 DEGs, with 2,132 upregulated genes, 2,201 downregulated genes, and 8,166 non-differentially expressed genes (**Fig. 3b; Suppl. Table 14**). Amongst the upregulated genes in line C3, we identified several expressed within the insect integument, including *tweedle* (*twdl*) *F*, *twdlG*, and *twdlV* as well as *cpr35B* and *cpr66D* ^32^, while several genes upregulated in line C4 are metabolic genes, including *cyp4d8, cyp6d4, hsp68,* and *gstS1* (**Fig. 3c; Suppl. Table 15**). This suggests that penetration resistance may confer spinosad resistance in line C3 while metabolic resistance may confer resistance in line C4. This also suggests that alleles resulting in either metabolic resistance or penetration resistance are present in the same field-collected spinosad-resistant *D. suzukii* population.

**Fig. 3:**
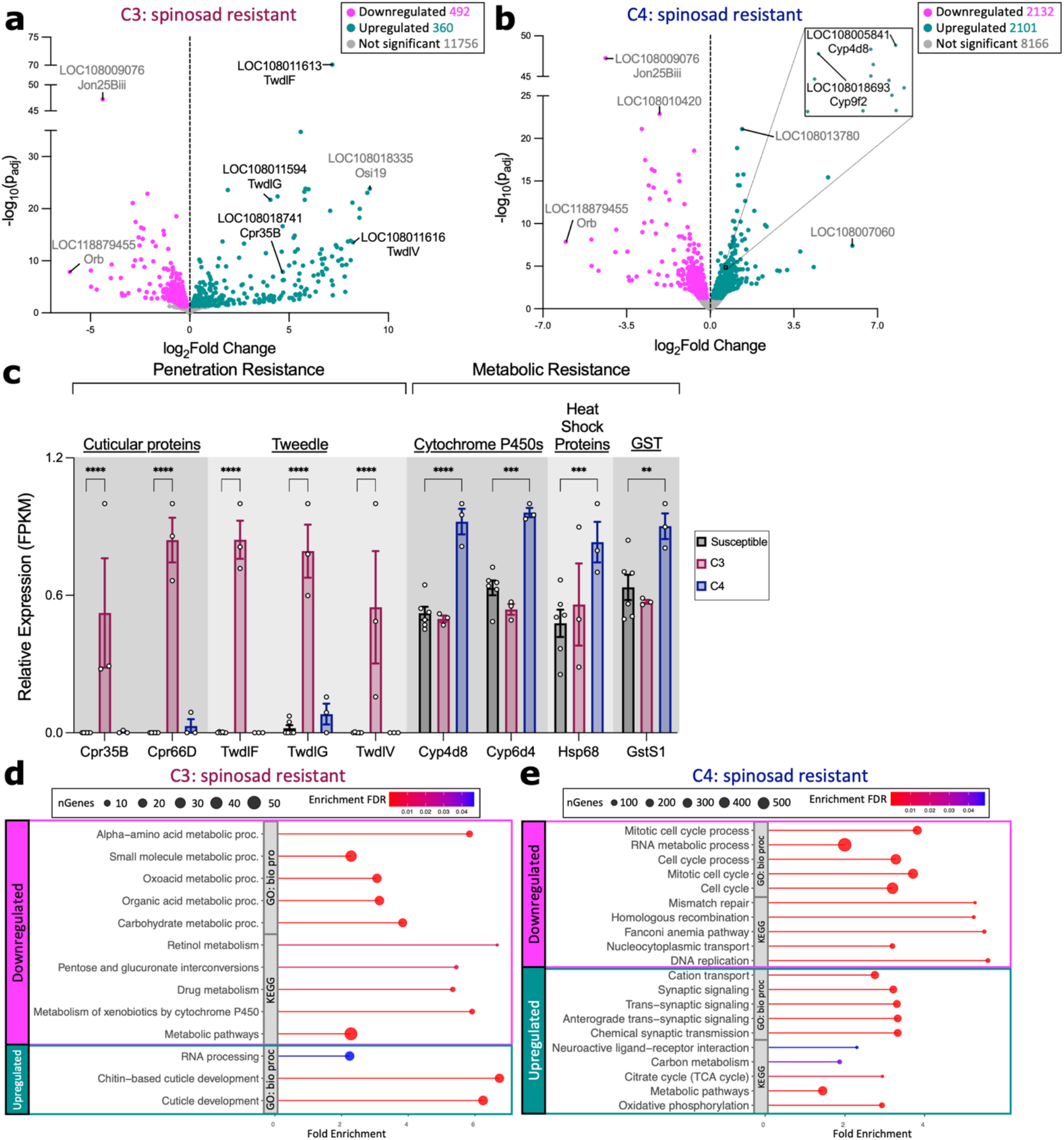
Spinosad-resistant lines exhibit increased expression of genes associated with either penetration resistance or metabolic resistance. (**a**) Volcano plot of genes displaying fold change gene expression differences between the resistant C3 line vs 2 susceptible lines (C2 and C5). Genes upregulated in the resistant populations (green) are to the right of the dotted line while genes downregulated in the resistant populations (pink) lie to the left of the dotted line. Genes that exhibit no significant difference (NS) in gene expression between the two populations are in grey. Highly significant differences are located higher up on the y-axis where p_adj_ is the Benjamin-Hochberg adjusted p-value. Labeled points signify genes that satisfy at least one of the following criteria: (1) have the largest fold change difference between the two groups, (2) have the lowest p_adj_, and (3) are genes known to be involved in insecticide resistance. Labels contain the *D. suzukii* gene symbol (LOC#########) and the corresponding *D. melanogaster* gene symbol. Genes known to be involved in conferring resistance are labeled in black while genes labeled in grey are not known to be directly involved in conferring resistance. (**b**) Volcano plot of genes displaying fold change gene expression differences between the resistant C4 line vs 2 susceptible lines (C2 and C5). The black box denotes the region zoomed-in in the insert. (**c**) Relative expression (FPKM) of metabolic and cuticular genes (*twdl*: *tweedle*; *cpr*: *cuticular protein*) in the susceptible (C2 and C5: black) and resistant (C3: maroon; C4: blue) groups extracted from the RNA sequencing data. Each point denotes a biological replicate (n=3 replicates of 8-10 females per line). Asterisks denote significant p-values as determined by 2-way ANOVA: **p<0.01, ***p<0.001, and ****p<0.0001. (**d**-**e**) Top 5 enrichment pathways within the Kyoto Encyclopedia of Genes and Genomes (KEGG) and Gene Ontology (GO) Biological Processes (bio proc) categories for genes up- or down-regulated in lines (**d**) C3 and (**e**) C4. The x-axis is Fold Enrichment, which is defined as the percentage of differentially expressed genes that belong to each pathway. Point size represents the number of genes (nGenes) within the category while color denotes the false discovery rate (FDR) correction of enrichment p-values.

Furthermore, we performed functional enrichment analyses and observed that genes downregulated in line C3 are enriched in metabolic pathways, including the metabolism of xenobiotics by the *cyp* pathway, while upregulated genes are enriched in pathways related to the insect cuticle and RNA processing (**Fig. 3d; Suppl. Table 16**). In line C4, on the other hand, genes downregulated in the resistant lines are enriched in pathways pertaining to cell cycle, DNA replication and repair, and cell division while upregulated genes are enriched in metabolism and neuronal signaling pathways (**Fig. 3e; Suppl. Table 17**).

WGCNA was performed to potentially identify novel gene clusters strongly correlated with spinosad resistance (**Suppl. Fig. 5-6**). In line C3, genes clustered into 47 different modules, with dark turquoise being most correlated with resistance (R^2^ = 0.83) (**Suppl. Fig. 5a; Suppl. Table 18**). Only 66 of the 79 genes within dark turquoise were functionally annotated and have a *D. melanogaster* homolog (**Suppl. Fig. 5b**). Since this module consists of such few genes, no genes were enriched in any pathways, however, there are a few genes in dark turquoise that are involved in chromatin organization, such as *histone H2A* ^33^, *modifier of mdg4* ^34^, and *histone methyl transferase 4-20* ^35^, as well as genes involved in hypoxia response (*ecdysone induced protein 93F* ^36^ and CG2918 ^37^) and negative regulation of cell growth (*La-related protein4B* ^38^ and *Forkhead box subunit O* ^39^). On the other hand, genes in line C4 were clustered into 42 colored modules, with green most correlated with resistance (R^2^ = 0.96) (**Suppl. Fig. 6a; Suppl. Table 19**). There are 371 genes in green, and of those, only 3 genes, *cyp6d4, cyp305A1,* and *GstO1,* belong to metabolic enzymes involved in insecticide detoxification ^24^ (**Suppl. Fig. 6b**). Green module genes are enriched in pathways involving neuronal organization and signaling as well as metabolism (**Suppl. Fig. 6c; Suppl. Table 20**). Therefore, genes most correlated with resistance in line C3 are genes that have not been previously implicated in conferring insecticide resistance in other insect species, whereas in line C4, 3 of the genes most correlated with resistance are within classes of metabolic enzymes known to promote insecticide resistance.

### New field-collected *Drosophila suzukii* populations in 2022 show evidence of increased metabolic resistance as compared to flies collected in 2019

With the identification of genes of interest that may confer insecticide resistance in isofemale lines of *D. suzukii*, we were interested in determining whether any of these genes are also differentially expressed in resistant *D. suzukii* recently collected from similar locations in California. We assessed the resistance status of the F1 of *D. suzukii* collected in 2022 from two strawberry fields using discriminating dose bioassays. As we showed previously, the mortality rates observed in the Wolfskill and susceptible isofemale lines were 100% in discriminating dose bioassays (**Fig. 1a-b**). In comparison, the mortality of F1 of both 2022 populations was significantly lower than 100% (**Fig. 4a**), suggesting that these populations are resistant to both zeta-cypermethrin and spinosad (zeta-cypermethrin: Population #1: t=23.88, df=9, p<0.0001; Population #2: t=20.82, df=9, p<0.0001) (spinosad: Population #1: t=6.736, df=9, p<0.0001; Population #2: t=6.708, df=9, p<0.0001).

**Fig. 4:**
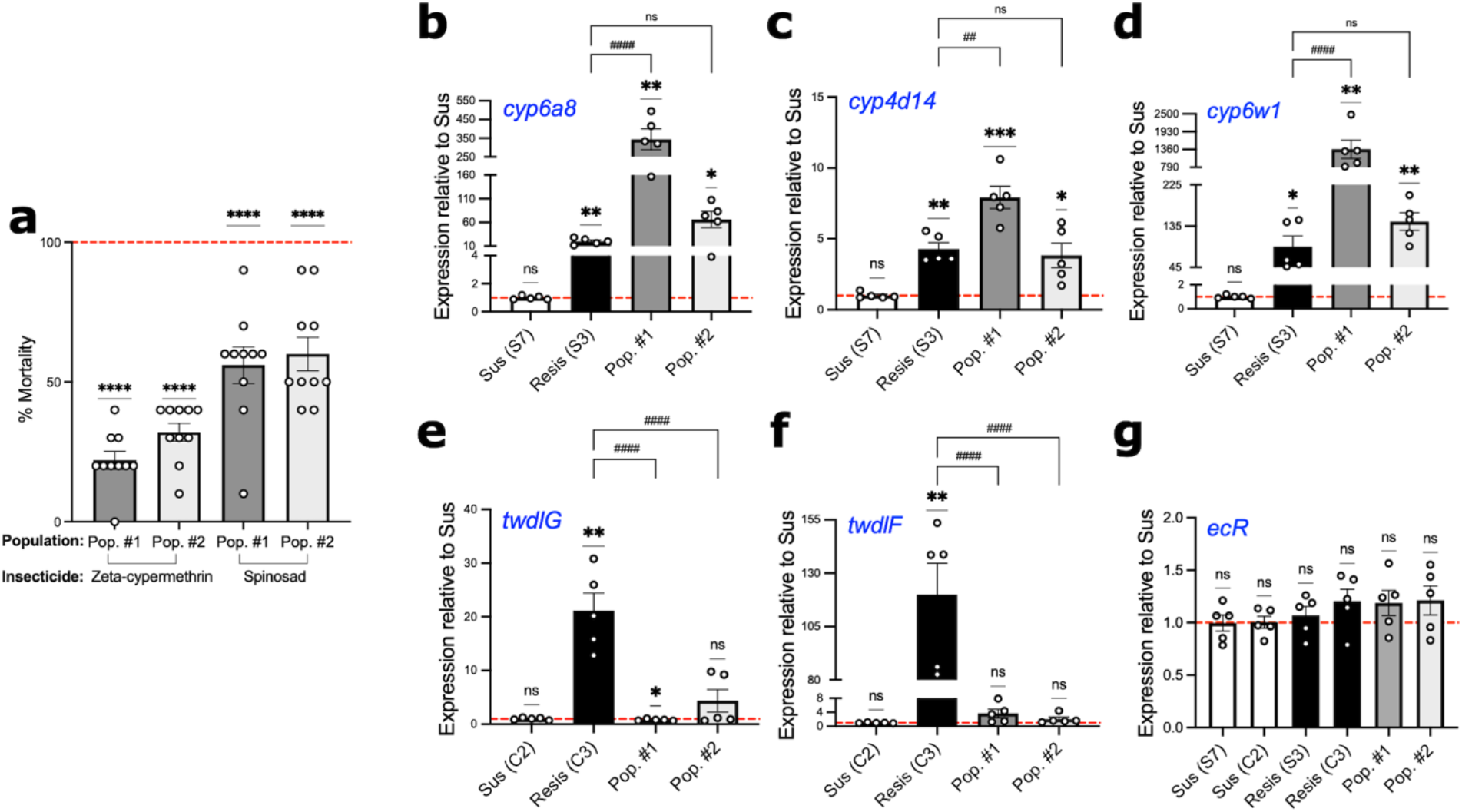
Field-collected *Drosophila suzukii* populations in 2022 show increased expression of metabolic genes that were differentially expressed in 2019 resistant populations. (**a**) Discriminating dose bioassay to assess mortality of 2022 field-collected *D. suzukii* populations (Pop. #1 and #2) when exposed to zeta-cypermethrin and spinosad. Each point represents a biological replicate of 5 males and 5 females (n=10), and error bars indicate ±SEM. Asterisks denote significant p-values as determined by one-sample t and Wilcoxon Test compared to a hypothetical mean of 100 (denoted by the dashed red line): **** p<0.0001. (**b**-**g**) Gene expression of (**b**-**d**) *cytochrome P450* (*cyp*) genes, (**e**-**f**) *tweedle* (*twdl*) genes, and (**g**) *ecdysone receptor* (*ecR*) in susceptible (S7 and C2) and resistant (S3 and C3) isofemale lines (established from 2019 collections) as well as 2022 field-collected populations (n=5 biological replicates of 8-10 females). Asterisks denote significant p-values as determined by one sample T and Wilcoxon Test compared to the average gene expression of the susceptible line (denoted by the red dashed line): **p<0.01 and **** p<0.0001. The hash marks (#) denote significant p-values as determined by One-way ANOVA followed by Holm-Sidak’s multiple comparisons test to assess expression differences between the 2022 F1 populations and the resistant line. Non-significant comparisons are denoted as “ns”.

Next, leveraging the results of our RNA-Seq experiment, we designed quantitative PCR (qPCR) primers to amplify five genes that were upregulated in at least one resistant *D. suzukii* isofemale line (**Fig. 4b-f**). Specifically, we detected *cyp6a8*, *cyp4d14*, and *cyp6w1* to evaluate metabolic resistance (**Fig. 4b-d**) and *twdlG* and *twdlF* to evaluate penetration resistance (**Fig. 4e-f**). We observed that both populations show increased expression of *cyp6a8* (**Fig. 4b; Suppl. Table 21**), *cyp4d14* (**Fig. 4c; Suppl. Table 21**), and *cyp6w1* (**Fig. 4d; Suppl. Table 21**) as compared to the susceptible controls. More so, the expression of all three *cyp* genes was significantly higher in Population #1 as compared to the resistant isofemale lines developed from 2019 field-collected populations (**Fig. 4b-d; Suppl. Table 21**). In fact, the expression of *cyp6a8* was 17.3-fold higher in Population #1 as compared to the isofemale resistant line from 2019 (**Fig. 4b**). Additionally, *cyp4d14* was 1.8-fold higher (**Fig. 4c**) while *cyp6w1* was 15.2-fold higher in Population #1 than in the 2019 resistant line (**Fig. 4d**).

We next assessed whether either of these lines exhibit penetration resistance by detecting cuticular genes *twdlG* (**Fig. 4e**) and *twdlF* (**Fig. 4f**). We observed a slightly lower expression of *twdlG* in Population #1 (**Fig. 4e**; **Suppl. Table 21**) and no significant difference of *twdlF* in either of the 2022 populations (**Fig. 4f**; **Suppl. Table 21**). Finally, we also detected expression of *ecdysone receptor* (*ecR*) as a negative control (**Fig. 4g**) because it was not previously observed to be differentially expressed in any of the isofemale resistant lines (**Suppl. Tables 4-5, 13-14**). There was no significant difference in *ecR* in either of the 2022 populations compared to the susceptible controls (**Fig. 4g**; **Suppl. Table 21**). Together, these results suggest that metabolic resistance confers insecticide resistance in the 2022 field-collected populations, as opposed to penetration resistance. Additionally, it shows that these qPCR-based assays are feasible and represents a more efficient approach of monitoring potential insecticide resistance in field-collected samples as opposed to performing bioassays to assess resistance.

To further confirm that the expression of these genes can reflect the susceptible vs resistance status of *D. suzukii*, we assayed the expression of *cyp6a8, cyp4d14, cyp6w1, twdlG,* and *twdlF* in two additional *D. suzukii* populations that were collected in Georgia, USA in 2023 and were found to be susceptible to zeta-cypermethrin and spinosad (**Suppl. Fig. 7f**), herein referred to as Populations #3 and #4. As expected, we observed that our pyrethroid-resistant line (S4) exhibited increased metabolic gene expression as compared to the susceptible Wolfskill line (S4: *cyp6a8* t=4.199, df=8, p=0.0179; *cyp4d14* t=7.577, df=8, p=0.0003; *cyp6w1* t=4.130, df=8, p=0.0193) (**Suppl. Fig. 7a-c**). The spinosad-resistant line (C3) also showed elevated expression in cuticular genes as compared to the susceptible Wolfskill line (C3: *twdlG* t=12.93, df=8, p<0.0001; *twdlF* t=5.691, df=8, p=0.0028) (**Suppl. Fig. 7d-e**). Populations #3 and #4, on the other hand, exhibited similar target gene expression levels as the susceptible Wolfskill control (Population #3: *cyp6a8* t=0.1687, df=8, p=0.9978; *cyp4d14* t=1.086, df=8, p=0.3090; *cyp6w1* t=0.01034, df=8, p=0.9978; *twdlG* t=0.7626, df=8, p=0.7165; *twdlF* t=0.1221, df=8, p=0.9992; and Population #4: *cyp6a8* t=0.003754, df=8, p=0.9978; *cyp4d14* t=2.036, df=8, p=0.1465; *cyp6w1* t=0.1595, df=8, p=0.9978; *twdlG* t=0.3463, df=8, p=0.7381; *twdlF* t=0.06005, df=8, p=0.992) (**Suppl. Fig. 7a-e**).

### Knockdown of *cyp4d14, cyp4d8*, and *cpr66D* increases susceptibility to insecticides

To determine whether the expression of metabolic and cuticular genes has a direct effect on insecticide resistance, we leveraged the genetic tools available in the closely related species *D. melanogaster* to manipulate the expression of these target genes and evaluate insecticide susceptibility using discriminating dose bioassays. We selected *D. melanogaster* mutant fly lines for three genes: *cyp4d14, cyp4d8,* and *cpr66D*. We chose a metabolic gene upregulated in both zeta-cypermethrin-resistant lines (S3 and S4), a gene associated with metabolic resistance in the spinosad-resistant line C4, and a gene implicated in penetration resistance that is upregulated in the spinosad-resistant line C3. We observed that reduced expression of *cyp4d14* (**Fig. 5a**; **Suppl. Table 22**) increases susceptibility to zeta-cypermethrin (**Fig. 5b**; **Suppl. Table 22**) and spinosad (**Fig. 5c**; **Suppl. Table 22**). However, we observed that reduced expression of *cyp4d8* (**Fig. 5d**; **Suppl. Table 22**) decreases susceptibility to zeta-cypermethrin (**Fig. 5e**; **Suppl. Table 22**) but increases susceptibility to spinosad (**Fig. 5f**; **Suppl. Table 22**). This is consistent with our differential expression analysis showing that *cyp4d8* is only upregulated in zeta-cypermethrin-resistant flies (**Fig. 2-3**). Finally, reduced expression of *cpr66D* (**Fig. 5g**; **Suppl. Table 22**) did not affect the susceptibility of flies to zeta-cypermethrin (**Fig. 5h**; **Suppl. Table 22**), but it increased susceptibility to spinosad (**Fig. 5i**; **Suppl. Table 22**). This result agrees with our sequencing analysis (**Fig. 2-3**), which showed that high levels of *cpr66D* was present in flies resistant to spinosad but not in flies resistant to zeta-cypermethrin.

**Fig. 5:**
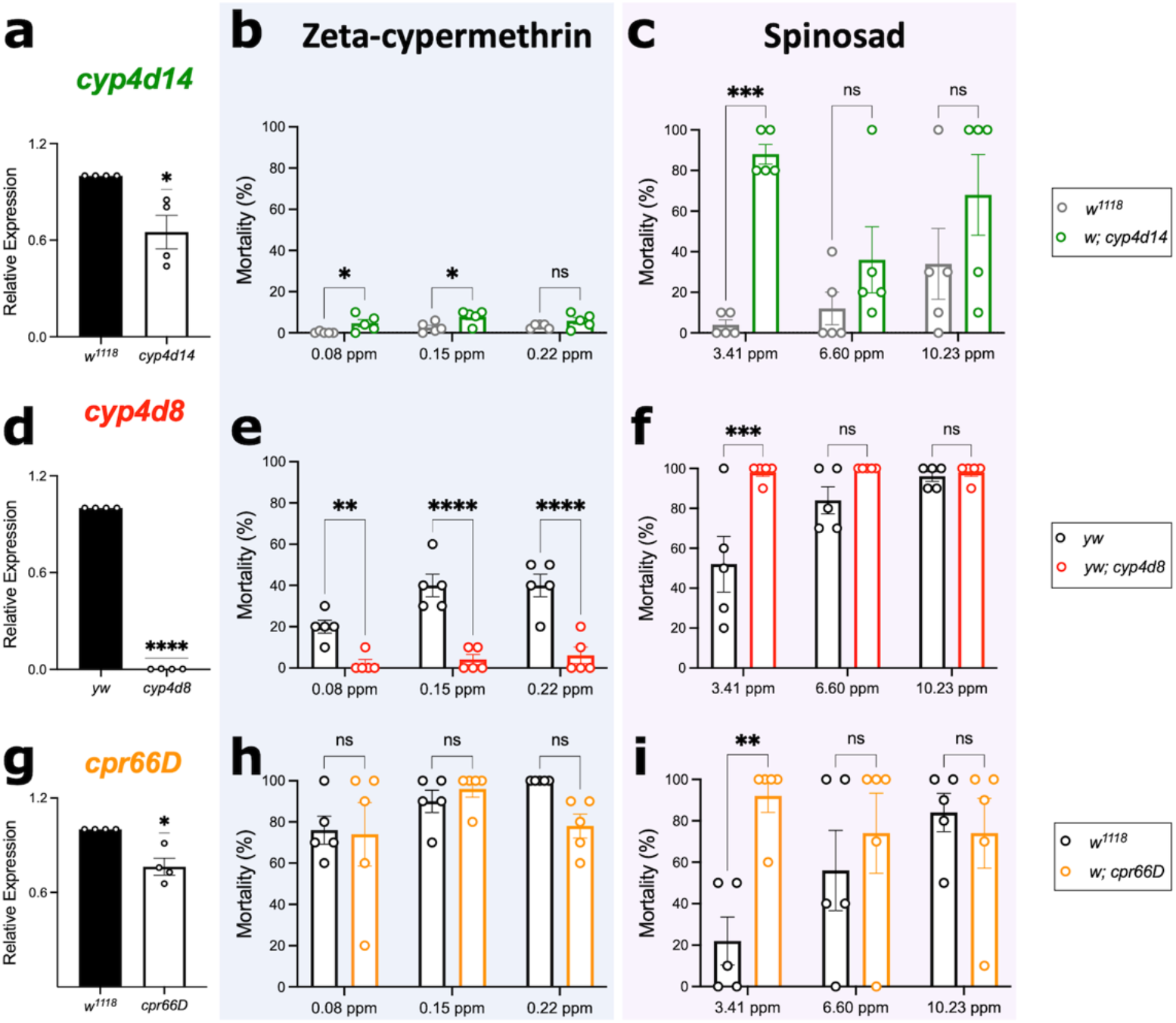
Reduced expression of *cyp4d14*, *cyp4d8*, and *cpr66D* affects *D. melanogaster* susceptibility to insecticides. (**a**) Relative expression of *cyp4d14* in wild-type (*w^1118^*) and *cyp4d14* mutant fly lines. Each point represents a biological replicate of 10 female bodies (n=4), and error bars indicate ±SEM. Asterisks denote significant p-values as determined by one-sample t and Wilcoxon Test compared to a hypothetical mean of 1: * p<0.05; **** p<0.0001. (**b**-**c**) Mortality of *w^1118^* (grey) and *cyp4d14* (green) flies when exposed to sublethal concentrations of (**b**) zeta-cypermethrin and (**c**) spinosad. Each point represents a biological replicate of 5 males and 5 females (n=5), and error bars indicate ±SEM. Asterisks denote significant p-values as determined by Two-way ANOVA followed by Holm-Sidak’s multiple comparisons test: * p<0.05; ** p<0.01; *** p<0.001; **** p<0.0001. Non-significant comparisons are denoted as “ns”. (**d**) Relative expression of *cyp4d8* in wild-type (*yw*) and *cyp4d8* mutant fly lines. (**e**-**f**) Mortality of *yw* (black) and *cyp4d8* (red) mutant flies when exposed to sublethal concentrations of (**e**) zeta-cypermethrin or (**f**) spinosad. (**g**) Relative expression of *cuticular protein* (*cpr*) *66D* in wild-type (*w^1118^*) or *cpr66D* mutant fly lines. (**h**-**i**) Mortality of *w^1118^* (grey) or *cpr66D* (orange) flies when exposed to sublethal concentrations of (**h**) zeta-cypermethrin or (**i**) spinosad.

### Sequence analysis reveals mutations in *para* or *nAChRα7* likely did not contribute to insecticide resistance

To investigate whether mutations within the target gene of each insecticide confer resistance, we assessed changes in allelic frequency between the resistant and susceptible populations of *D. suzukii* (**Suppl. Fig. 8a-c**). Within the gene that encodes the target of zeta-cypermethrin, the VGSC *para*, we identified a significant difference in allelic frequency in line S3 at nucleotide position 3658093, which is located within intron 30 in the gene body, and one difference in allelic frequency at nucleotide position 3655811, which is located within intron 29 in the gene body, in line S4 (**Suppl. Fig. 8b-c**; 3658093: S3 p= 0.005171, S4 p=0.6199; 3655811: S3 p=0.4725, S4 p=0.001131). Although we did not identify a mutation within the protein-coding region of *para*, we did observe that *para* is downregulated in zeta-cypermethrin-resistant *D. suzukii* (**Fig. 2a-b**). Therefore, it is possible that reduced expression of the target protein, rather than a site-specific mutation, contributes to resistance in these flies. This mechanism has not been previously evaluated in the context of pyrethroid resistance.

We then analyzed the spinosad target protein *nAChRα7,* the *D. suzukii* homolog of *D. melanogaster nAChRα6* inferred by sequence similarity. We identified three nucleotide positions within the 5’ untranslated region (UTR) that exhibit a significant change in allelic frequency in line C3 and no changes in line C4 (**Suppl. Fig. 8d-g**; 6579905: C3 p=0.0101, C4 p=0.06667; 6580106: C3 p=0.01515, C4 p=0.2424; 6580194: C3 p=0.04762, C4 p=1). Given we did not identify a mutation within the protein-coding region of the gene or any differential expression of *nAChRα7* (**Suppl. Table 14-15**) in the resistant lines, we suspect that target-site resistance is not an underlying mechanism for spinosad resistance in these lines. However, we cannot rule out that the allelic frequency changes we observed in the 5’ UTR may affect *nAChRα7* protein levels given that the 5’UTR is important for translation initiation (reviewed in ^40^).

## Discussion

Insecticide resistance in the invasive agricultural pest *D. suzukii* has been detected in California, U.S.A. over the past several years, but the molecular mechanisms driving these changes have yet to be identified ^20–22^. We developed isofemale lines from field-collected populations of *D. suzukii* resistant to either zeta-cypermethrin or spinosad to identify the molecular mechanisms underlying insecticide resistance. We sequenced the transcriptomes of two resistant lines per population and found evidence of metabolic resistance in zeta-cypermethrin-resistant *D. suzukii* (**Fig. 6**). Specifically, we observed an upregulation of genes encoding metabolic detoxification enzymes in zeta-cypermethrin-resistant *D. suzukii*. Interestingly, we also observed decreased expression of the target gene *para*. This does not constitute conventional target-site resistance as resistant lines do not have a mutation in the target gene, but rather, an overall decrease in the target gene could render the insecticide less effective. This mechanism can be tested in a future mechanistic study. In *D. suzukii* resistant to spinosad, we identified evidence of penetration resistance in one line (C3), reflected by the upregulation of several genes expressed in the insect cuticle such as *tweedle* genes and *cuticle proteins* (*cpr*). In the other resistant line (C4) however, we observed evidence of metabolic resistance reflected by an upregulation of metabolic genes. Our results for the spinosad-resistant lines reveal the possibility for multiple mechanisms of insecticide resistance to be present in a single population.

**Fig. 6:**
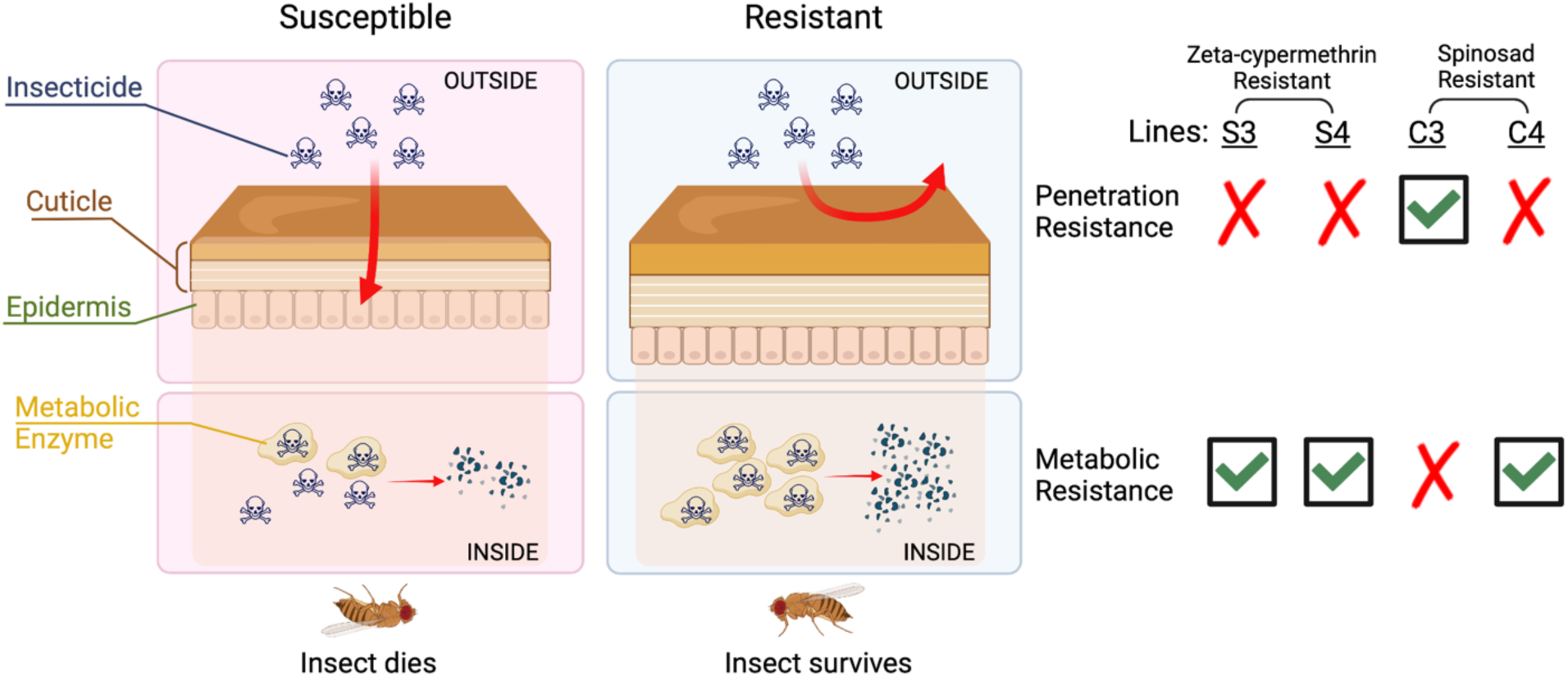
Schematic representation of the molecular mechanisms conferring either zeta-cypermethrin or spinosad-resistance in *Drosophila suzukii*. The cuticle of susceptible *D. suzukii* is permeable to insecticides, enabling the insecticide to enter the insect and bind to its target protein, ultimately killing the insect. However, in the case of zeta-cypermethrin-resistant *D. suzukii*, an increased expression of metabolic enzymes results in an increased breakdown of the insecticide before it can bind to its target protein. Spinosad-resistant *D. suzukii* have increased expression of cuticular genes such that the cuticle is less penetrable by insecticides, allowing them to survive. Additionally, spinosad-resistant *D. suzukii* can also exhibit an upregulation of metabolic enzymes to increase detoxification of the insecticide, promoting the survival of the flies. This figure was created with BioRender.com (license to laboratory of JCC).

We concluded from our differential gene expression (DEG) analysis and weighted gene co-expression network analysis (WGCNA) that metabolic resistance and penetration resistance are contributing to pyrethroid and spinosad resistance observed in *D. suzukii* in California. However, it is important to note that we cannot rule out that other mechanisms are also contributing to the observed resistance, given that we detected many other differentially expressed genes and our WGCNA focused on the module with the highest correlation to resistance. Our DEG analysis uncovered other potential genes and pathways that may be important in *D.suzukii* resistance development, and future experiments will be necessary to explore the role of these differentially expressed genes. For example, we observed that genes involved in RNA processing and splicing are enriched in differentially expressed genes in zeta-cypermethrin and spinosad resistant lines (**Fig. 2d**; **Fig. 3d**). Splicing is a biological process that produces proteins with diverse structures and functions encoded by a singular gene (reviewed in ^41^). Therefore, it is possible that resistant isofemale lines may undergo differential splicing, resulting in widespread differences in gene and isoform expression as compared to susceptible flies. It has been shown in other insect species that differential expression of various isoforms of *nAChRα6* confers insecticide resistance ^42, 43^, but at present it is not clear whether changes in splicing are limited to specific genes or observed more broadly in the transcriptome. Results from this study set the stage for future studies into other potential mechanisms of insecticide resistance, including the possibility of alternative splicing as a driver for resistance.

We leveraged our findings to design molecular diagnostics, specifically quantitative PCR (qPCR) assays, that could identify insecticide resistance in the field (**Fig. 4**). Thus far, insecticide resistance in *D. suzukii* has only been detected in California ^20–22^. Therefore, our diagnostic tests can be used to monitor insecticide resistance development in California and can be used to detect early development of resistance in locations where resistance has yet to be reported. This would allow growers to adjust spray programs and delay and/or prevent resistance development in the fly population. Utilizing a few genes that were differentially expressed between the resistant and susceptible lines, such as *cytochrome P450* (*cyp*) and *tweedle* genes, we designed qPCR assays to monitor resistance development. A benefit to using molecular diagnostics to detect resistance, as opposed to insecticide bioassays, is that they require few individuals (as little as 5 flies) as input whereas bioassays require a much larger number of flies. A similar molecular diagnostic detecting *cyp* expression to identify metabolic resistance has been previously developed and validated in mosquitoes ^44^. More so, beyond just validating our diagnostic, we observed significantly higher levels of *cyp* expression in a 2022 field-collected population (**Fig. 4**), suggesting that resistance has not only persisted but increased since the 2019 collection. This observation is consistent with the general trend of increased spinosad resistance in *D. suzukii* from 2018 to 2020 ^20^.

Additionally, we observed that reduced expression of a single *cyp* or *cpr* gene that we found to be differentially expressed in resistant *D. suzukii* renders *D. melanogaster* more susceptible to insecticides (**Fig. 5**), supporting the hypothesis that increased expression of metabolic and cuticular genes promote resistance. It is important to note this experiment leveraged the genetic mutants available in the closely related species *D. melanogaster,* as opposed to creating transgenic *D. suzukii*, which is much more challenging. Thus, it is possible that knocking down expression of other genes we identified to be differentially expressed in the resistant lines may not have the desired effect, if any, on susceptibility. This is because the proteins involved in insecticide resistance may differ, even between species of the same genus. For example, the substrate specificity of the various *cyps* may vary from species to species ^45^. We also observed that the effect of gene knockdown is more evident at lower concentrations of insecticide. Furthermore, we cannot rule out that altering the expression of one *cyp* gene does not affect the expression of other *cyp* genes. This may explain why the *cyp4d8* mutant exhibits increased susceptibility to spinosad but decreased susceptibility to zeta-cypermethrin (**Fig. 5e-f**). Finally, it is possible that knocking down several metabolic genes simultaneously will produce a stronger phenotype, given that several metabolic genes are differentially expressed in our dataset. This may be because multiple *cyps* may target the same substrate ^45^.

Our study also provides insights into the possibility of cross-resistance. Currently, in order to delay the development of resistance, there are restrictions on how many applications of a single type of insecticide are permitted in a site ^46^. As a result, organic berry growers alternate usage between other organically-approved insecticides and spinosads ^12, 47, 48^. Therefore, it is important to understand whether the mechanisms conferring resistance to one insecticide enables the insect to be resistant to other classes of insecticides as well. For instance, because the spinosad-resistant line C3 has increased expression of cuticular genes (**Fig. 3c**) suggesting a less penetrable cuticle, it is possible that the integument is less penetrable to other insecticides as well, but that remains to be tested. Furthermore, zeta-cypermethrin-resistant *D. suzukii* and spinosad-resistant line C4 express high levels of many metabolic enzymes implicated in metabolic resistance (**Fig. 2c** and **3c**). There are studies in house flies and onion thrips that attribute spinosad resistance to an upregulation in *cyp* expression ^49, 50^. Notably, our 2022 collections of *D. suzukii* exhibit resistance to both zeta-cypermethrin and spinosad and appear to be resistant through an upregulation of metabolic genes (**Fig. 4**), and knockdown of *cyp4d14* increase both zeta-cypermethrin and spinosad susceptibility (**Fig. 5b-c**). Thus, our results suggest a high likelihood for cross-resistance. Additional experiments are needed to verify this prediction for cross-resistance.

Finally, we anticipate the ability to leverage our results to optimize current *D. suzukii* management practices. For instance, we identified an upregulation of metabolic enzymes in resistant *D. suzukii* (**Fig. 2** and **Fig. 3c**). Presumably, this upregulation will increase detoxification, rendering the insecticide less effective. To combat this effect, synergists can be applied in conjunction with insecticides. Synergists are metabolic enzyme inhibitors, so when used in conjunction with insecticides, synergists can increase insecticide efficacy ^51^. An alternative approach would be for growers to adopt an integrated pest management (IPM) strategy to control *D. suzukii*. IPM promotes increased control of a pest by adopting a combination of different strategies including genetic, biological, cultural, and chemical control ^52^. It is possible that alternating between the use of insecticides and the use of non-chemical methods of control will alleviate the pressure driving insecticide resistance such that it is selected out of the population. Based on our data, we speculate that there is a high fitness cost associated with insecticide resistance. For example, in zeta-cypermethrin-resistant flies, we observed differential expression of many genes involved in neuronal system development and signaling (**Fig 2c**), suggesting that neuronal processing is affected, potentially compromising a wide range of behaviors such as mating and feeding ^53, 54^. In the case of spinosad-resistant flies, we observed that many of the downregulated genes are enriched with metabolic pathways, suggesting that spinosad-resistant flies have energy usage deficiencies. It is possible that the fitness costs associated with resistance are the reason that it took eleven years for insecticide resistance to develop since the invasion of *D. suzukii* into California despite intense spray programs ^46^. Further experiments will need to be conducted to identify the costs associated with resistance in *D. suzukii.* Moreover, it is possible that in the absence of selective pressure caused by spraying insecticides, in combination with the fast generation time of *D. suzukii* and their short lifespans ^46^, alternating between insecticide spraying and other forms of pest control can be more effective at controlling *D. suzukii* infestations in the field. In fact, a previous study ^20^ has demonstrated that resistance increases throughout the growing season, likely due to increased exposure to insecticides from multiple applications. Therefore, it is possible that a short-term halt in spraying of insecticides of a specific chemistry for a few generations would increase susceptibility, given that the selective pressure is removed. Experiments are currently in progress to assess how long resistance persists in a population after spraying has ceased. Further, to combat penetration resistance, insecticides can be administered in bait traps as opposed to spraying such that the insecticide enters the flies through the digestive system rather than through the insect cuticle.

In conclusion, our study characterizes the mechanisms of insecticide resistance in *D. suzukii* collected in California, U.S.A. We provide evidence that metabolic and penetration resistance underlie insecticide resistance in the populations we sampled from. Additionally, we developed and validated molecular assays that can monitor resistance in field populations of *D. suzukii*. Finally, our study provides insights into the possibility of cross-resistance and information that can be used to improve *D. suzukii* management programs.

## Methods

### Field *Drosophila suzukii* populations and development of isofemale lines

To assess resistance to zeta-cypermethrin (Mustang® Maxx 0.8 EC, FMC Corporation, Philadelphia, PA), isofemale lines were established from *D. suzukii* adults reared from fruits collected in October 2019 from a strawberry field in Monterey County, CA, U.S.A. Sixty fruits were collected and transported to the laboratory of Dr. Frank Zalom at the University of California, Davis. Twenty of the fruits were transferred to a plastic container containing a layer of cotton topped with sand as a substrate for pupation, for a total of three containers. The containers were maintained at 23 ± 1°C, 55-65% relative humidity (RH), and a 14-hour light:10-hour dark photoperiod in a walk-in growth chamber (Percival Scientific Inc., Perry, IA) and checked daily until fly emergence. Emerged *D. suzukii* flies were separated from non-target species and reared in bottles containing Bloomington standard *Drosophila* cornmeal diet (https://bdsc.indiana.edu/information/recipes/bloomfood.html).

To assess resistance to spinosad (Entrust® SC 22.5% spinosad, a mixture of spinosad A & D, Corteva Agriscience, Indianapolis, IN), isofemale lines were established from *D. suzukii* adults collected from a caneberry field in Santa Cruz County, CA, U.S.A. in November 2019. Adult flies were live-captured using McPhail traps (Great Lakes IPM, Inc., Vestaburg, MI) baited with approximately 20 ml of a yeast (7 g)-sugar (113 g)-water (355 ml) solution. Traps were collected the next day and returned to the laboratory. Flies were anesthetized using CO_2_ to facilitate the removal of any non-target species (approximately twenty females and twenty males per bottle) and transferred into diet bottles (described above).

Field-collected *D. suzukii* were assessed for resistance as described below in the “Discriminating dose bioassays” section. Each isofemale line was established from a single wild-caught, gravid, non-insecticide-treated female from a resistant population ^55^. Crossing of siblings was repeated for eight generations for each isofemale line. A total of eight isofemale lines were established for each site, for a total of sixteen lines. Bioassays were performed once isofemale lines were established to identify resistant and susceptible isofemale lines.

Field-collected populations (referred to as Populations #1 and #2) used for quantitative PCR (qPCR) assays were reared from fruits collected from two California open strawberry fields in Santa Cruz County in September 2022. A hundred ripe fruits were collected from each field and placed in plastic containers and transported to the laboratory. Fruits were transferred to new plastic containers containing a layer of cotton topped with sand. For each site, five containers of twenty fruits were prepared and placed at 23±1°C, 55-65% RH, 14-hour light: 10-hour dark photoperiod and checked daily until fly emergence. Emerged flies were aspirated into diet bottles. Twenty female and twenty male *D. suzukii* adults were then moved to new diet bottles for propagation to increase total available flies, and the progeny (F1) from each site were used in bioassays. The susceptible field-collected line, Population #3, was collected from blueberries in Alma, GA, U.S.A. in June 2023 while Population #4 was also collected from blueberries in Broxton, GA, U.S.A. in July 2023. Flies were maintained on standard diet food and sent to the laboratory of Dr. Joanna Chiu at UC Davis for use in molecular assays.

### Drosophila melanogaster strains

*Drosophila melanogaster* mutant fly lines for genes demonstrated to be upregulated in resistant *D. suzukii* were obtained from the Bloomington *Drosophila* Stock Center (BDSC) (Bloomington, IN): *w^11^*^18^*; cytochrome P450 cyp4d14* (stock no. 24262: *Mi{GFP[E.3xP3]=ET1}Cyp4d14[MB03260] w^1118^), yw; cyp4d8* (stock no. 58019: *y[1] w[*]; Mi{y[+mDint2]=MIC}Cyp4d8[MI13043]*), and *w^1118^; cuticular protein (cpr) 66D* (stock no. 17735: *w^1118^; PBac{w[+mC]=PB}Cpr66D[c05937])* ^56–58^. These mutant fly lines have transgenes inserted into an intronic region in either *cyp4d14, cyp4d8,* and *cpr66d* (**Suppl. Fig. 9**). Because not all transgene insertions in intronic regions will affect gene expression, we confirmed that the mutants have decreased expression of inserted genes using qPCR (**Fig. 5a, 5d,** and **5g**).

### Discriminating dose bioassays

To identify resistant isofemale lines, discriminating dose bioassays were performed using a glass vial residue bioassay. Briefly, the interior of 20-ml glass scintillation vials (Fisher Scientific, Pittsburgh, PA) were coated with insecticide solution at the discriminating dose ^15, 59^. Excess insecticide was removed, and treated vials and caps were placed upright in a fume hood to dry overnight. Five male and five female adults (3-5 days after emergence) from each isofemale line were transferred into each treated vial. The vials were then maintained at 23±1°C, 55-65% RH, 14-hour light: 10-hour dark photoperiod for the duration of the experiment.

Susceptibility of the established isofemale lines to zeta-cypermethrin (Mustang® Maxx 0.8 EC, FMC Corporation, Philadelphia, PA) was assessed at the discriminating dose of 6.89 mg/liter (which is eight times the concentration required to kill 90% of the tested population) ^22, 60^. Based on mortality at this concentration, two resistant lines and one susceptible line were selected per site for dose-response bioassays.

Susceptibility of isofemale lines to spinosad (Entrust® SC 22.5% spinosad, a mixture of spinosad A & D, Corteva Agriscience, Indianapolis, IN) was assessed at the discriminating dose of 928 mg/liter ^20, 21^. The insecticide was diluted in Induce (Helena Chemical Company, Memphis, TN) – deionized water solution at a rate of 1266 μl Induce per liter of deionized water. Induce is a non-ionic surfactant used to help spread the solution uniformly and increase adherence to the vial surfaces. Two resistant lines and one susceptible isofemale line were selected for dose-response bioassays.

Mortality was assessed after 6 hours of zeta-cypermethrin exposure and after 8 hours of spinosad exposure. Moribund and dead individuals were combined and considered as dead. Moribund flies are defined as those showing clear signs of toxicity (i.e. slow uneven movements, leg twitching, and inability to hold on to the bioassay vial). Despite being alive, these flies would not recover from the insecticide. Differences of mortality between isofemale lines and the Wolfskill control were assessed by one-way ANOVA followed by Tukey’s multiple comparison test using GraphPad Prism Version 9.3.1 (GraphPad Software, La Jolla, CA).

Discriminating dose bioassays performed in *D. melanogaster* were performed as described above but with the following modifications. Five replicates consisting of 5 males and 5 females were tested at 3 doses of each insecticide: 0.08ppm, 0.15ppm, and 0.22ppm for Mustang® Maxx and 3.41ppm, 6.00ppm, and 10.23ppm Entrust. Differences of mortality between the wild-type control and the mutant were assessed by two-way ANOVA followed by Holm-Sidak’s multiple comparison test using GraphPad Prism.

### Dose-response bioassays

To identify the lethal concentration required to kill 50% of the tested population (LC_50_), various concentrations of 1-18 mg/liter zeta-cypermethrin (0ppm, 1ppm, 2ppm, 4ppm, 6.89ppm, 10ppm, 12ppm, 15ppm, and 18ppm) were tested for the resistant isofemale lines. For the susceptible isofemale line, six concentrations of 0.2-10 mg/liter zeta-cypermethrin (0ppm, 0.2ppm, 0.5ppm, 1ppm, 2ppm, 4ppm, 6.89ppm) were tested. Mortality was recorded after six hours, and the number of dead flies was recorded. For the resistant isofemale lines, several concentrations of 100-10,000 mg/liter spinosad (100ppm, 300ppm, 928ppm, 3000ppm, 6000ppm, 8000ppm, and 10,000ppm) were tested while six concentrations of 30-3,000 mg/liter spinosad (30ppm, 65ppm, 100ppm, 300ppm, 928ppm, and 3000ppm) were tested for the susceptible isofemale line. Eight replicates were tested per concentration, and mortality of flies was recorded after eight hours of spinosad exposure.

Mortality data from dose-response bioassays were fitted to a two-parameter log-logistic model with a lower limit of 0 and an upper limit of 1 using the ‘drc’ package^61^ in R 4.2.1 ^62^. The function ‘ED’ was used to calculate estimated effective doses (LC_50_ values). Pairwise z-tests were conducted with the ‘compParm’ function to compare LC_50_ values between different isofemale lines to further analyze susceptibility. All models included insecticide concentration (mg/liter) and isofemale line as predictor variables, while the response variable was the proportion of dead flies. Proportions were weighted by total sample size in each vial. As a control, we included 2 isofemale lines established from flies collected in an untreated orchard located near Winters in Solano County, CA ^21^, referred as Wolfskill populations.

### RNA extraction, library preparation, and high throughput sequencing

Female *D. suzukii* flies were entrained at 25°C in 12-hour light:12-hour dark cycles for two full days. On the third day, flies were collected on dry ice sixteen hours after lights-on. This time point was selected because *D. suzukii* was previously observed to exhibit a low level of *cytochrome P450* expression at this time ^63^. This means any overexpression may be more easily observed. Fly bodies were separated from heads using frozen metal sieves (Newark Wire Cloth Company, Clifton, NJ). Eight to ten female bodies were homogenized using a Kimble^®^ pellet pestle^®^ Cordless Motor (DWK Life Sciences, Rockwood, TN) and Kimble™ Kontes™ Pellet Pestle™ (DWK Life Sciences) in 300 μL TRI reagent (Sigma, St. Louis, MO). 60 μL of 100% chloroform (Sigma) was added and incubated at room temperature for 10 minutes. The upper aqueous layer was recovered after spinning down and transferred to a new microcentrifuge tube. RNA was precipitated with an equal volume of 100% isopropanol at −20°C overnight. After spinning down, the RNA pellet was washed with 70% ethanol once and allowed to air dry. The pellet was then resuspended in 20 μL 1X Turbo DNA-free kit buffer (Thermo Fisher Scientific, Waltham, MA) and treated with Turbo DNase per manufacturer’s instructions. RNA quality was assessed with both the Agilent 2100 Bioanalyzer system (Agilent Technologies, Santa Clara, CA) and the Qubit RNA IQ kit (Invitrogen, Waltham, MA) on the Qubit 4 Fluorometer (Invitrogen). RNA purity was measured with the Nanodrop 1000 (Thermo Fisher Scientific), and RNA quantity was measured with the Qubit RNA HS (high sensitivity) assay kit (Invitrogen) on the Qubit 4 Fluorometer.

Illumina short-read sequencing libraries were prepared with 1 μg of high-quality RNA and the TruSeq Stranded mRNA Library Prep Kit (Illumina, San Diego, CA) according to manufacturer’s protocol. A total of twenty-four libraries were prepared: three biological replicates of two zeta-cypermethrin-resistant lines, two zeta-cypermethrin-susceptible lines, two spinosad-resistant lines, and two spinosad-susceptible lines. Library insert size and quality was measured with the Agilent 2100 Bioanalyzer System. Library concentration was measured with the Qubit 4 Fluorometer. All libraries generated from zeta-cypermethrin-resistant and susceptible *D. suzukii* lines were pooled together, and all libraries generated from spinosad-resistant and susceptible *D. suzukii* lines were pooled together, such that there were twelve libraries per pool. Pooled samples were sent to Novogene (Sacramento, CA) for sequencing on the HiSeq X Ten platform (Illumina) using PE150.

### Differential gene expression analysis

Differential gene expression analysis was performed using sequencing reads derived from Illumina short-read sequencing. First, rRNA reads were removed using SortMeRNA v2.1 ^64^. Adapters (ILLUMINACLIP parameters 2:30:10) and low-quality ends (LEADING: 10, TRAILING:10, MINLEN:36) were trimmed using Trimmomatic v0.35 ^65^. Cleaned reads were aligned to the NCBI *Drosophila suzukii* Annotation Release 102 based on the LBDM_Dsuz_2.1.pri assembly (accession no. GCF_013340165.1) ^66^ using STAR v2.7.9a ^67^. Count data from STAR (--quantMode GeneCounts) served as input in the DESeq2 package ^68^ in R to perform differential expression analysis on each resistant line vs both susceptible samples. Each resistant line was compared to susceptible samples separately as each line might exhibit resistance due to different mechanisms. Genes with fold change differences between resistant vs susceptible populations with a Benjamini-Hochberg adjusted p-value < 0.05 were considered differentially expressed. Expression levels of genes were also measured as fragments per kilobase of exon per million mapped (FPKM) values calculated with Stringtie v2.0.4 ^69^. The consistency between biological replicates was calculated with Pearson’s correlation coefficient, which was determined with the ‘stats’ package in R version 4.2.1. Expression differences of key genes between the resistant and susceptible populations were calculated with two-way ANOVA followed by two-stage linear set-up procedure of Benjamini, Krieger, and Yekutieli on GraphPad Prism.

### Weighted Gene Co-expression Network Analysis

Gene expression (in FPKM) served as input for Weighted Gene Co-expression Network Analysis (WGCNA). Genes with an expression value of zero for more than six samples were excluded from analysis. To explore the modules most correlated with insecticide resistance, a correlation analysis using resistance status was performed with the WGCNA package (Version 1.72.1) ^31^ on R. Modules with a p-value < 0.05 were considered significant. Functional enrichment analysis (described below) was performed on the module with the highest correlation with resistance.

### Functional enrichment analysis

Genes were functionally annotated using BLAST+ (Version 2.12.0) against the NCBI *Drosophila melanogaster* Annotation r6.32 based on the Release 6 plus ISO1 mitochondrial genome assembly (accession no. GCA_000001215.4) ^70^. Gene Ontology (GO) enrichment of genes were performed using ShinyGO 0.76.3 ^71^. GO terms and pathways were considered enriched if the false discovery rate (FDR) < 0.05.

### Variant calling

To identify allelic changes between the resistant and susceptible populations in the target genes *paralytic* and *nicotinic acetylcholine receptor subunit alpha 7*, variants were called using Freebayes v1.3.5 ^72^. Differences between allelic counts between the resistant and susceptible groups were compared using Fisher’s Exact Test on R v4.2.1.

### Quantitative polymerase chain reaction (qPCR) for gene expression analysis

Total RNA extraction from F3 of the 2022 collection (see “Field *Drosophila suzukii* populations and development of isofemale lines”) was performed as described above (see “RNA extraction, library preparation, and high throughput sequencing”). cDNA synthesis was performed with the SuperScript IV Reverse Transcriptase kit (ThermoFisher Scientific) according to the manufacturer’s instructions, using 4 μg of RNA as input. cDNA was then diluted ten-fold with nuclease-free water. qPCR was performed using Sso Advanced SYBR green supermix (Bio-Rad, Hercules, CA) in a CFX384 (Bio-Rad). Primer sequences are listed in **Suppl. Table 23**. Cycling conditions were 95°C for 30 seconds followed by forty cycles of 95°C for 5 seconds and an annealing/extension phase at 60°C for 30 seconds. The reaction was concluded with a melt curve analysis from 65°C to 95°C in 0.5°C increments at 5 seconds per step. Three technical replicates were performed per biological replicate for a total of five biological replicates. Isofemale lines served as the susceptible and resistant controls. Resistant control lines were selected based off whether the line overexpressed the gene of interest in the differential gene expression analysis. Data were analyzed using the standard ΔΔCt method ^73^, and target gene mRNA expression levels were normalized to the reference gene, *ribosomal protein L32* (*rpL32*) ^74^. Finally, relative expression was calculated by dividing the expression level for each sample by the average expression of the susceptible control for all biological replicates such that the expression of the susceptible line would be one. To compare the expression levels of each sample to the susceptible control, a one sample t and Wilcoxon test were performed on GraphPad Prism. Additionally, gene expression levels of the resistant control line were compared to the expression level of the 2022 populations using one-way ANOVA followed by Holm-Sidak’s multiple comparisons test on GraphPad Prism. Experiments using *D. melanogaster* were similarly performed but with the following modifications. Three technical replicates were performed per biological replicate for a total of four biological replicates. The wild-type control was determined based on the genetic background of the mutant fly (see “*Drosophila melanogaster* strains”). Targeted gene expression was normalized to the reference gene *cbp20* ^75^. To compare the expression levels of each mutant line to the wild-type control, a one sample t and Wilcoxon test were performed with a hypothetical mean of 1 using GraphPad Prism.

## Acknowledgements

We thank the Bloomington *Drosophila* Stock Center for providing *D. melanogaster* stocks. We also thank all members of the Chiu laboratory and members of the SCRI spotted wing *drosophila* research team, in particular Rufus Isaacs, Steven Van Timmeren, Elizabeth Beers, Gregory Loeb, and Vaughn Walton for their feedback and suggestions. This project is supported by the National Institute of Food and Agriculture at the United States Department of Agriculture, awards 2020-51181-32140 and CA-D-ENM-2150-H to JCC, and the CDFA Specialty Crop Block Grant 21SCBPCA1002 awarded to JCC.

## Author contributions

CAT, JCC, and FGZ conceived the study; FGZ, FG, CJ, and AAS conducted field collections of fly populations; CAT, FG, CCT, and NLN performed bioassays, and CAT and FG performed accompanying statistical analyses; CAT and CCT performed RNA extraction for sequencing, and CAT prepared Illumina sequencing libraries; CAT, CRC, KML, SH, and JCC performed bioinformatic analyses; CAT and CHC designed primers; CAT conducted molecular genetics experiments; CAT, FG, FZ, and JCC contributed to critical interpretation of the data; CAT wrote the paper with the input of JCC and all other authors.

## Data Availability Statement

Raw Illumina reads have been deposited to the NCBI Sequence Read Archive (SRA) and can be found under BioProject accession number PRJNA983428. Supplementary Tables 4-5, 7-14, and 16-20 contain the lists of differentially expressed genes, the results of the functional enrichment analyses, and WGCNA and have been uploaded to Dryad (https://datadryad.org/stash/share/J1eywW3gpjEL2FzQIUOsshzg3hnnr_GHX-IKv7Qw2DM).

