## Supplemental Figures and Tables for "Transcriptome analysis of *Drosophila suzukii* reveals molecular mechanisms conferring pyrethroid and spinosad resistance"

### Supporting information:

**Supplemental Figs. 1 to 9 and Supplemental Tables 3 and 23**

**Note that the following are provided as .xlsx files on Dryad**
(https://datadryad.org/stash/share/J1eywW3gpjEL2FzQIUOsshzg3hnnr_GHX-IKv7Qw2DM)

**Suppl. Table 1:** Statistics for zeta-cypermethrin bioassays
**Suppl. Table 2**: Statistics for spinosad bioassays
**Suppl. Table 4**: DEGs in zeta-cypermethrin-resistant *Drosophila suzukii* isofemale line (line S3) relative to susceptible lines (lines S7 and S8) developed from the same field-collected population
**Suppl. Table 5**: DEGs in zeta-cypermethrin-resistant *Drosophila suzukii* isofemale line (line S4) relative to susceptible lines (lines S7 and S8) developed from the same field-collected population
**Suppl. Table 6**: Statistics comparing the expression of metabolic genes in zeta-cypermethrin-resistant vs. susceptible *Drosophila suzukii*
**Suppl. Table 7**: Enrichment of DEGs in zeta-cypermethrin-resistant *Drosophila suzukii* (line S3)
**Suppl. Table 8**: Enrichment of DEGs in zeta-cypermethrin-resistant *Drosophila suzukii* (line S4)
**Suppl. Table 9**: Genes in all WGCNA modules for zeta-cypermethrin-resistant *Drosophila suzukii* line S3
**Suppl. Table 10**: Enrichment of the turquoise module in zeta-cypermethrin-resistant *Drosophila suzukii* line S3
**Suppl. Table 11**: Genes in all WGCNA modules for zeta-cypermethrin-resistant *Drosophila suzukii* line S4

**Suppl. Table 12**: Enrichment of the turquoise module in zeta-cypermethrin-resistant *Drosophila suzukii* line S4
**Suppl. Table 13**: DEGs in spinosad-resistant *Drosophila suzukii* isofemale line (line C3) relative to susceptible lines (lines C2 and C5) developed from the same field-collected population
**Suppl. Table 14**: DEGs in spinosad-resistant *Drosophila suzukii* isofemale line (line C4) relative to susceptible lines (lines C2 and C5) developed from the same field-collected population
**Suppl. Table 15**: Statistics comparing the expression of metabolic and cuticular genes in spinosad-resistant vs. susceptible *Drosophila suzukii*
**Suppl. Table 16**: Enrichment of DEGs in spinosad-resistant *Drosophila suzukii* (line C3)
**Suppl. Table 17**: Enrichment of DEGs in spinosad-resistant *Drosophila suzukii* (line C4)
**Suppl. Table 18**: Genes in all WGCNA modules for spinosad-resistant *Drosophila suzukii* line C3
**Suppl. Table 19**: Genes in all WGCNA modules for spinosad-resistant *Drosophila suzukii* line C4
**Suppl. Table 20**: Enrichment of the green module in spinosad-resistant *Drosophila suzukii* line C4
**Suppl. Table 21**: Statistics for bioassays with 2022 *Drosophila suzukii* populations
**Suppl. Table 22**: Statistics for bioassays with *Drosophila melanogaster* mutants

| | | Resistant | | | | | | Susceptible | | | | | |
|---|---|---|---|---|---|---|---|---|---|---|---|---|---|
| | | **S3_1** | **S3_2** | **S3_3** | **S4_1** | **S4_2** | **S4_3** | **S7_1** | **S7_2** | **S7_3** | **S8_1** | **S8_2** | **S8_3** |
| Resistant | **S3_1** | 1.000 | | | | | | | | | | | |
| | **S3_2** | 0.968 | 1.000 | | | | | | | | | | |
| | **S3_3** | 0.973 | 0.976 | 1.000 | | | | | | | | | |
| | **S4_1** | 0.943 | 0.937 | 0.957 | 1.000 | 0.975 | 0.964 | | | | | | |
| | **S4_2** | 0.954 | 0.956 | 0.957 | 0.975 | 1.000 | 0.988 | | | | | | |
| | **S4_3** | 0.943 | 0.956 | 0.950 | 0.964 | 0.988 | 1.000 | | | | | | |
| Susceptible | **S7_1** | 0.804 | 0.860 | 0.828 | 0.759 | 0.806 | 0.823 | 1.000 | 0.984 | 0.973 | | | |
| | **S7_2** | 0.854 | 0.901 | 0.880 | 0.825 | 0.868 | 0.874 | 0.984 | 1.000 | 0.975 | | | |
| | **S7_3** | 0.803 | 0.861 | 0.846 | 0.779 | 0.810 | 0.824 | 0.973 | 0.975 | 1.000 | | | |
| | **S8_1** | 0.929 | 0.957 | 0.954 | 0.938 | 0.954 | 0.958 | 0.877 | 0.914 | 0.880 | 1.000 | 0.986 | 0.981 |
| | **S8_2** | 0.917 | 0.950 | 0.946 | 0.917 | 0.936 | 0.947 | 0.893 | 0.923 | 0.900 | 0.986 | 1.000 | 0.989 |
| | **S8_3** | 0.926 | 0.958 | 0.957 | 0.919 | 0.938 | 0.951 | 0.879 | 0.912 | 0.900 | 0.981 | 0.989 | 1.000 |

**Value**
1.000
0.750

**Suppl. Fig. 1: Heatmap of Pearson's correlation coefficients between zeta-cypermethrin-resistant and susceptible lines.** Comparisons between lines resistant (S3, S4) or susceptible (S7, S8) to zeta-cypermethrin. The numbers following the underscore indicate biological replicates (n=3), each consisting of 8-10 female flies. Comparisons between biological replicates are indicated by black boxes. Highly correlated samples are in blue while less correlated samples are in red.

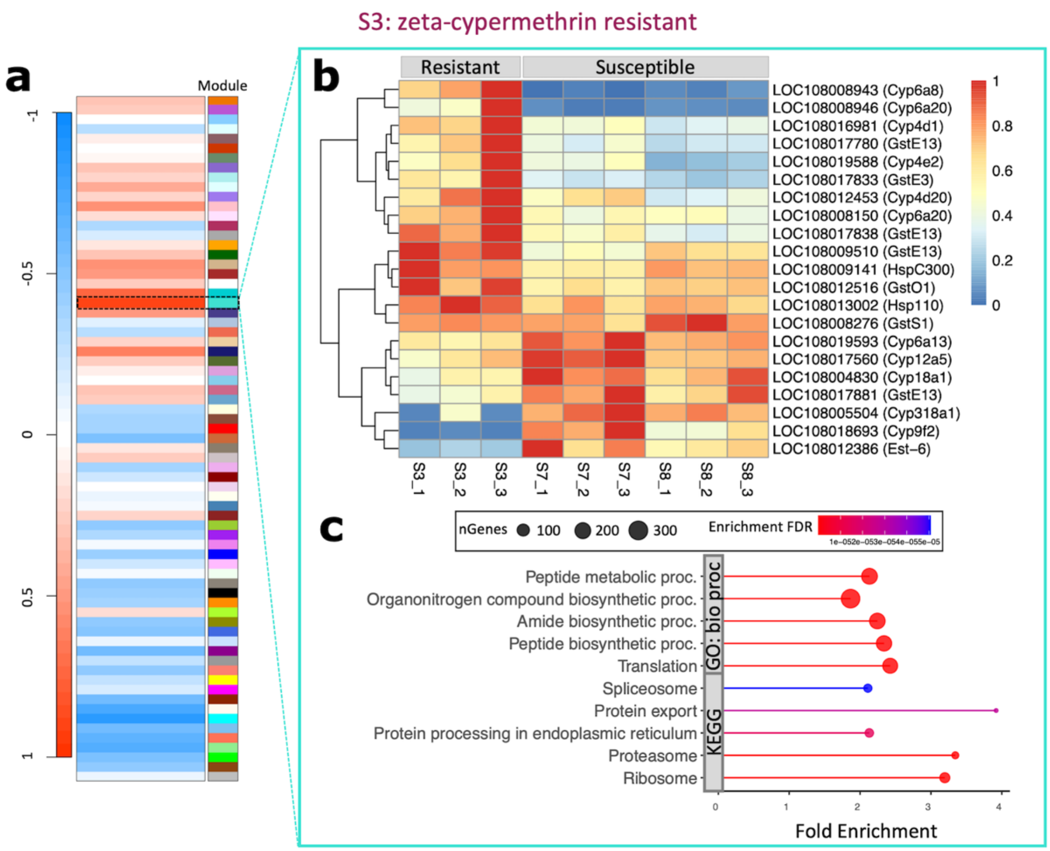


**Suppl. Fig. 2: Genes involved in metabolic resistance are highly correlated with zeta-cypermethrin resistance in *Drosophila suzukii* line S3.** (**a**) Heat map of gene clusters determined by Weighted Gene Correlation Network Analysis (WGCNA). Modules that are positively correlated with resistance are red while those that are negatively correlated are blue. (**b**) Heat map showing the expression of metabolic genes (in FPKM) within the turquoise module. Labels contain the *D. suzukii* gene symbol (LOC#########) and the corresponding *D. melanogaster* gene symbol. Red indicates high expression while blue indicates low expression. Line S3 is resistant while lines S7 and S8 are susceptible. The number following the underscore indicates different biological replicates. (**c**) Top 5 enrichment pathways within the Kyoto Encyclopedia of Genes and Genomes (KEGG) and Gene Ontology (GO) Biological Processes (bio proc) categories for genes within the turquoise module. The x-axis is Fold Enrichment, which is defined as the percentage of differentially expressed genes that belong to each pathway. Point size represents the number of genes (nGenes) within the category while color denotes the false discovery rate (FDR) correction of enrichment p-values.

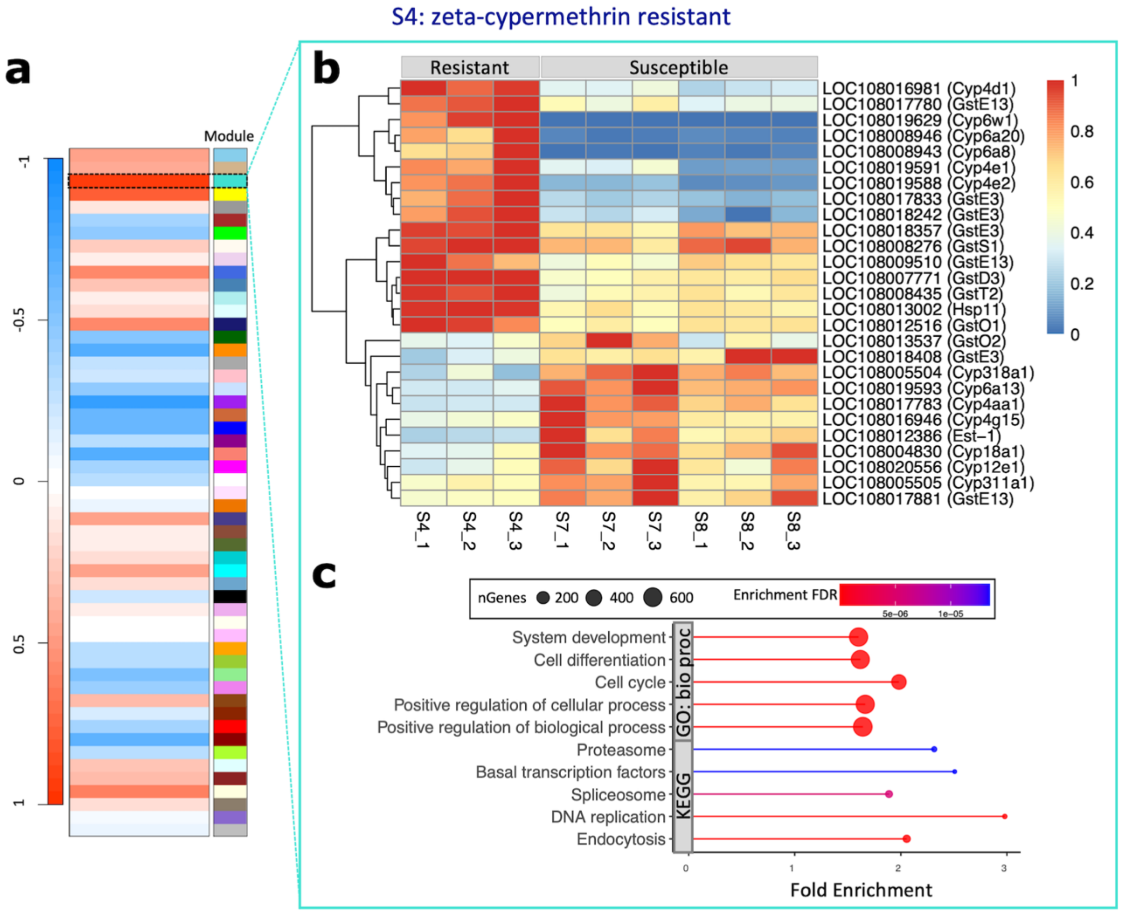


**Suppl. Fig. 3: Genes involved in metabolic resistance are highly correlated with zeta-cypermethrin resistance in *Drosophila suzukii* line S4.** (**a**) Heat map of gene clusters determined by Weighted Gene Correlation Network Analysis (WGCNA). Modules that are positively correlated with resistance are red while those that are negatively correlated are blue. (**b**) Heat map showing the expression of metabolic genes (in FPKM) within the turquoise module. Labels contain the *D. suzukii* gene symbol (LOC#########) and the corresponding *D. melanogaster* gene symbol. Red indicates high expression while blue indicates low expression. Line S4 is resistant while lines S7 and S8 are susceptible. The number following the underscore indicates different biological replicates. (**c**) Top 5 enrichment pathways within the Kyoto Encyclopedia of Genes and Genomes (KEGG) and Gene Ontology (GO) Biological Processes (bio proc) categories for genes within the turquoise module. The x-axis is Fold Enrichment, which is defined as the percentage of differentially expressed genes that belong to each pathway. Point size represents the number of genes (nGenes) within the category while color denotes the false discovery rate (FDR) correction of enrichment p-values.

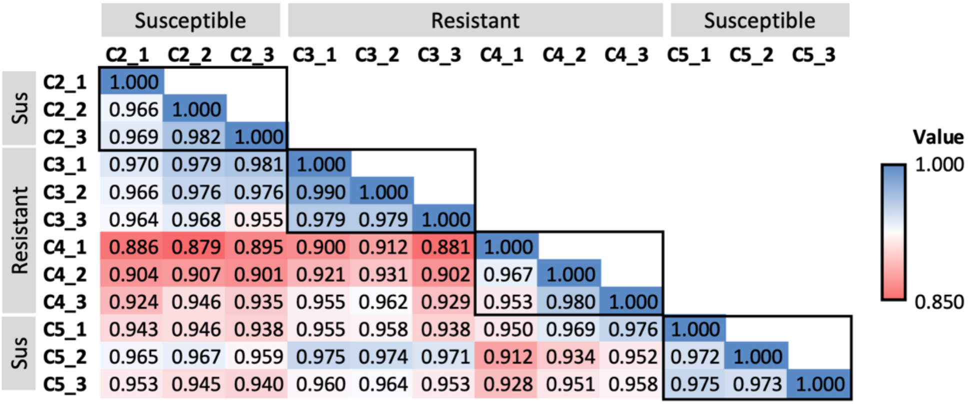


**Suppl. Fig. 4: Heatmap of Pearson's correlation coefficients between spinosad-resistant and susceptible lines.** Comparisons between lines resistant (C3, C4) or susceptible (C2, C5) to spinosad. The numbers following the underscore indicate biological replicates (n=3). Comparisons between biological replicates are indicated by black boxes. Highly correlated samples are in blue while less correlated samples are in red.

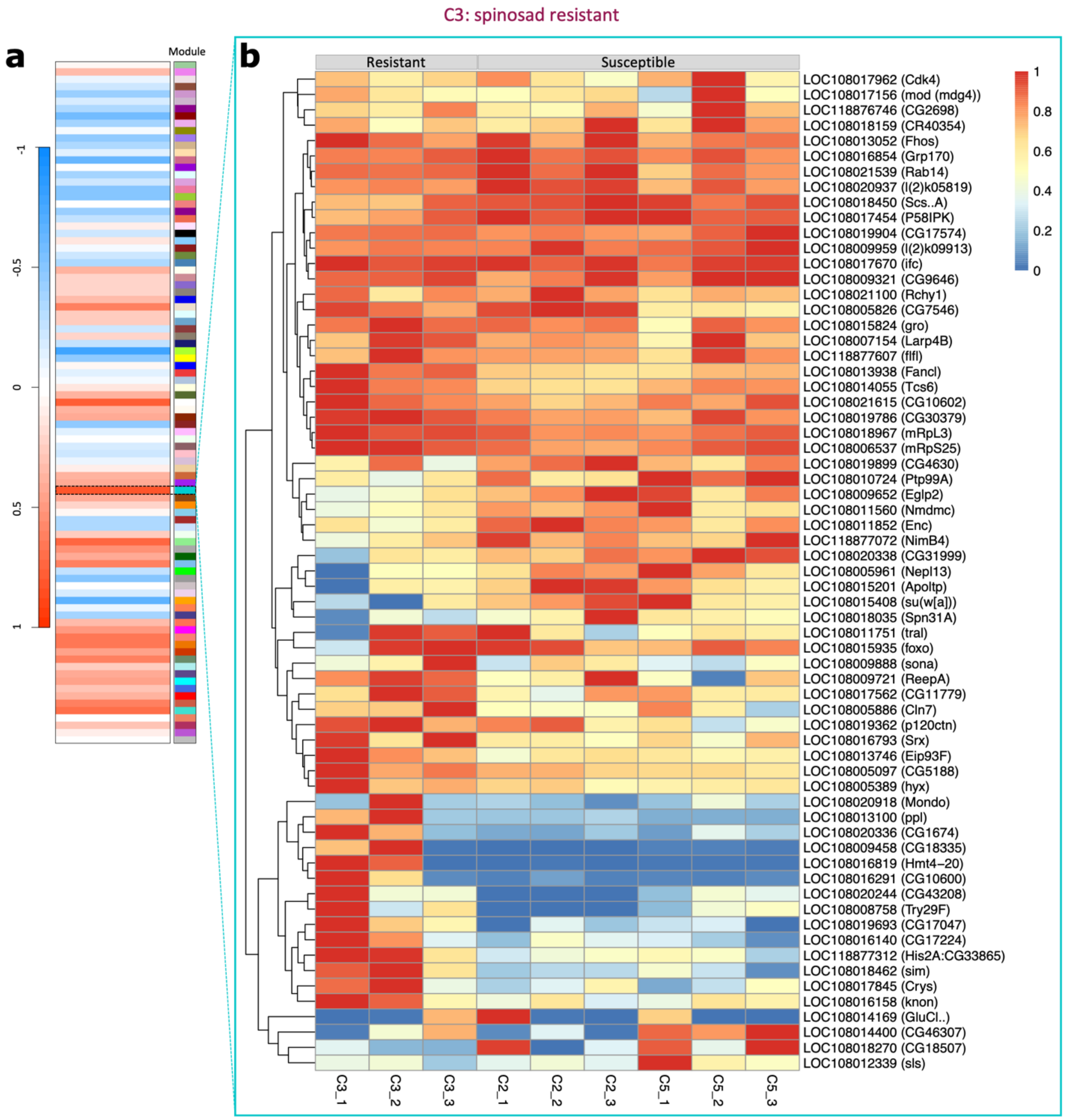


**Suppl. Fig. 5: Genes highly correlated with spinosad resistance in *Drosophila suzukii* line C3.** (**a**) Heat map of gene clusters determined by Weighted Gene Correlation Network Analysis (WGCNA). Modules that are positively correlated with resistance are red while those that are negatively correlated are blue. (**b**) Heat map showing the expression of all genes (in FPKM) within the dark turquoise module. Labels contain the *D. suzukii* gene symbol (LOC#########) and the corresponding *D. melanogaster* gene symbol. Red indicates high expression while blue indicates low expression. Line C3 is resistant while lines C2 and C5 are susceptible. The number following the underscore indicates different biological replicates.

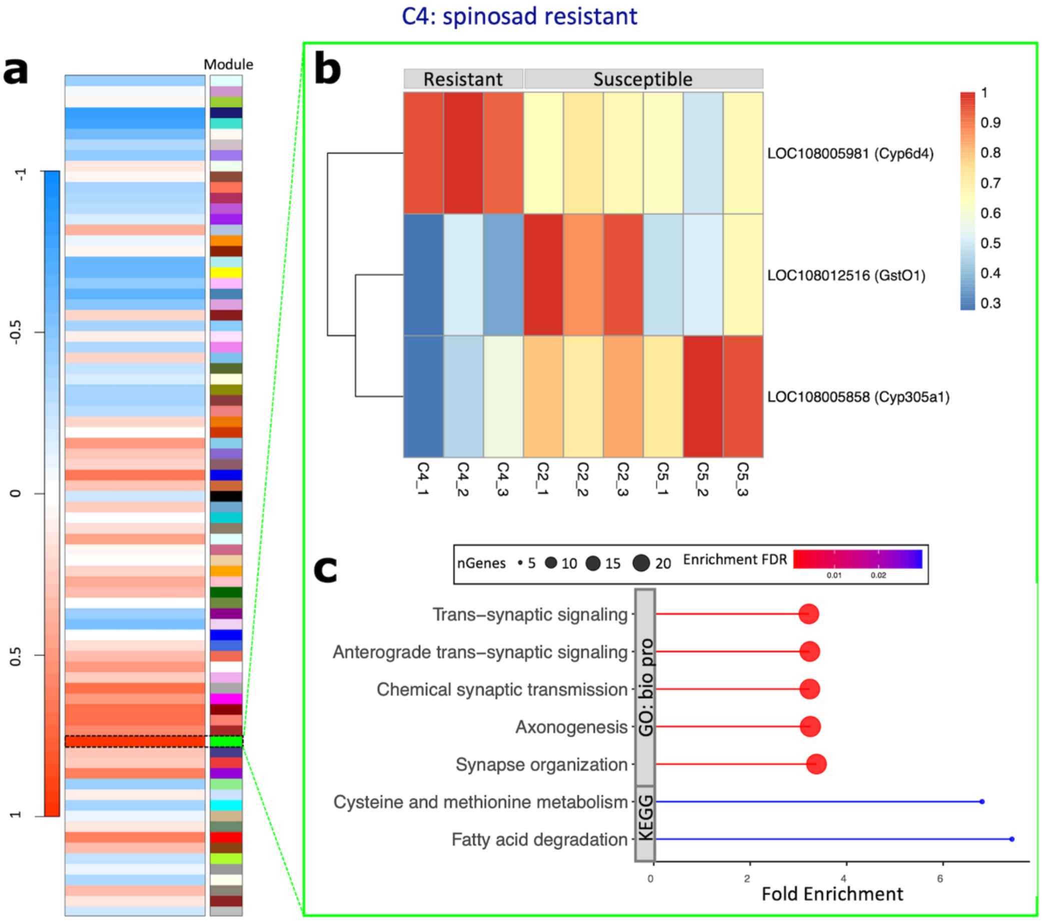


**Suppl. Fig. 6: Genes highly correlated with spinosad resistance in *Drosophila suzukii* line C4 include metabolic genes.** (**a**) Heat map of gene clusters determined by Weighted Gene Correlation Network Analysis (WGCNA). Modules that are positively correlated with resistance are red while those that are negatively correlated are blue. (**b**) Heat map showing the expression of metabolic genes (in FPKM) within the green module. Labels contain the *D. suzukii* gene symbol (LOC#########) and the corresponding *D. melanogaster* gene symbol. Red indicates high expression while blue indicates low expression. Line C4 is resistant while lines C2 and C5 are susceptible. The number following the underscore indicates different biological replicates. (**c**) Enriched pathways within the Kyoto Encyclopedia of Genes and Genomes (KEGG) category and the top 5 pathways within Gene Ontology (GO) Biological Processes (bio proc) category for genes within the green module. The x-axis is Fold Enrichment, which is defined as the percentage of differentially expressed genes that belong to each pathway. Point size represents the number of genes (nGenes) within the category while color denotes the false discovery rate (FDR) correction of enrichment p-values.

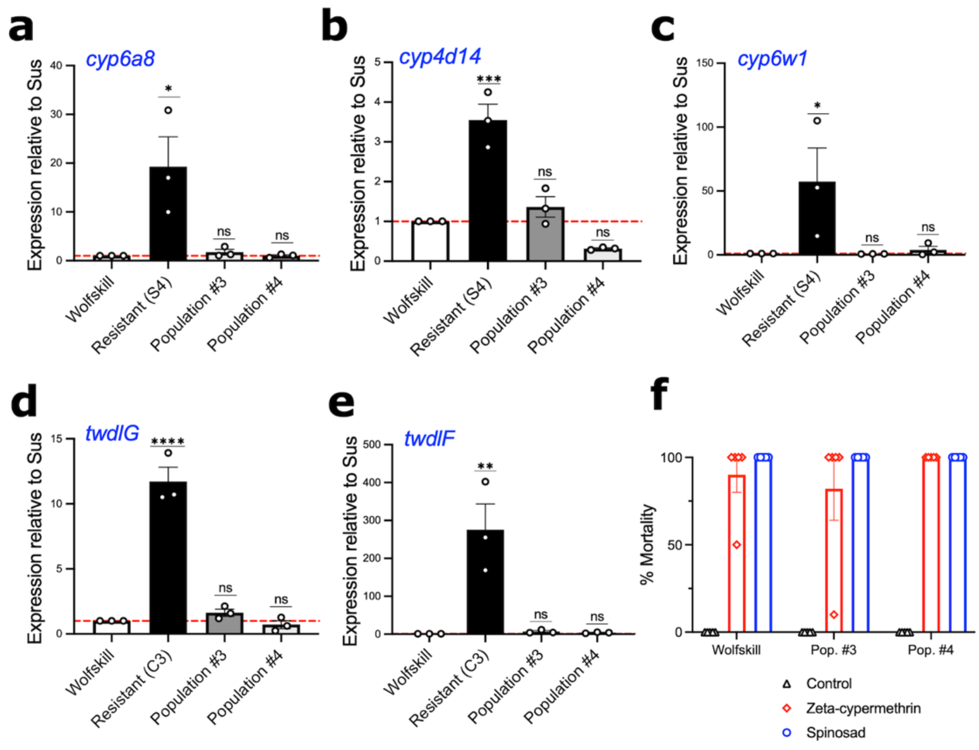


**Suppl. Fig. 7: Susceptible *D. suzukii* populations collected in 2023 in Georgia, USA, express similar levels of *cytochrome P450* genes and *tweedle* genes as the Wolfskill susceptible control.** (**a**-**e**) Gene expression of (**a**-**c**) *cytochrome P450* (*cyp*) genes and (**d**-**e**) *tweedle* (*twdl*) genes in susceptible *D. suzukii* populations collected in either California, U.S.A. (Wolfskill) or Georgia, USA (Populations #3 and #4) and resistant isofemale lines (S4 and C3) established from 2019 California collections (n=3 biological replicates of 5 females). Asterisks denote significant p-values as determined by One-way ANOVA followed by Holm-Sidak's multiple comparisons test when compared to the gene expression of the Wolfskill susceptible line (denoted by the red dashed line): *p<0.05, **p<0.01, ***p<0.001, and **** p<0.0001. Non-significant comparisons are denoted as "ns". (**f**) Discriminating dose bioassay to assess mortality of field-collected *D. suzukii* populations (Pop. #3 and #4) when exposed to either zeta-cypermethrin (red; diamonds) or spinosad (blue; circles). Each point represents a biological replicate of 5 males and 5 females (n=5). Control vials consisted of no insecticide (black; triangles).

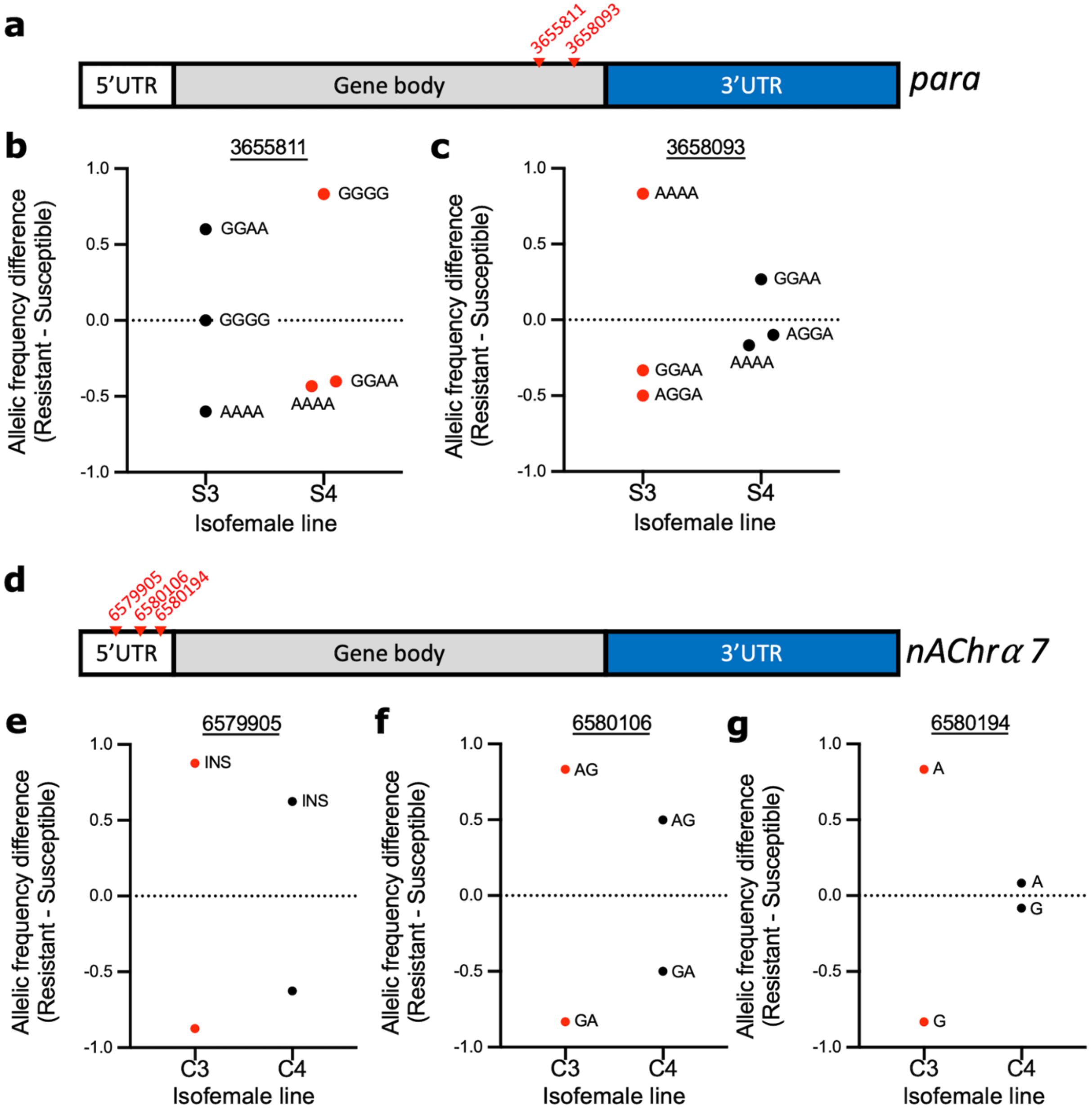


**Suppl. Fig. 8: Insecticide-resistant lines exhibit allele frequency differences in the insecticide target protein.** (**a**) Schematic of the gene, *paralytic* (*para*), the target protein of zeta-cypermethrin. Red labels indicate genomic loci shown in panels **b** and **c**. (**b-c**) Scatter plots of allele frequency differences on chromosome NW_023496846.1 at positions (**b**) 3655811 and (**c**) 3658093, which are both located within introns in the gene body of *para,* between each zeta-cypermethrin resistant line (S3 or S4; x-axis) vs both susceptible lines (S7 and S8). Each point is labeled with the allele variant. Red points indicate significant allele frequency differences between at least one resistant line and both susceptible lines as determined by Fisher's Exact Test while black points indicate differences that are not significantly different between the resistant and susceptible lines. Alleles located above the dotted line are more prevalent in the resistant line while alleles that fall below the line are more prevalent in the susceptible lines. (**d**) Schematic of the gene, *nicotinic acetylcholine receptor alpha* 7 (*nAChr*$\alpha$*7)*, the target of spinosad. The red labels indicate positions on chromosome NW_023496800.1 shown in panels **e**-**g**. (**e-g**) Scatter plots of allele frequency differences at positions (**e**) 6579905, (**f**) 6580106, and (**g**) 6580194, which are located within exons in the gene body of *nAchr*$\alpha$*7,* between each spinosad-resistant line (C3 or C4) vs both susceptible lines (C2 and C5). Insertions are labeled as "INS."

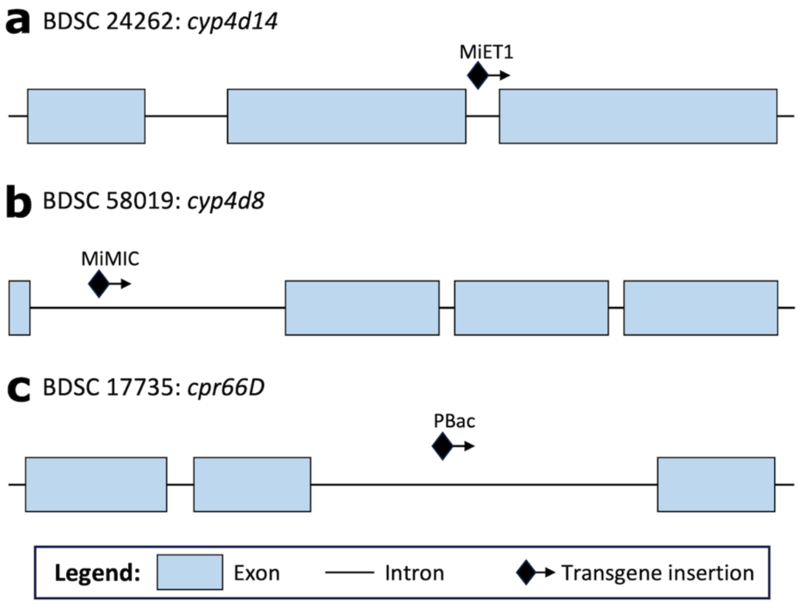


**Suppl. Fig. 9: Diagram of transgenic insertions for *Drosophila melanogaster* flies used in discriminating dose bioassays and quantitative PCR assays, Related to Figure 5.** *D. melanogaster* fly lines expressing an inserted transgene in (**a**) *cyp4d14* (BDSC stock no. 24262) (**b**) *cyp4d8* (BDSC stock no. 58019), and (**c**) *cpr66D* (BDSC stock no. 17735), where MiET1 (Minos element), MiMIC (Minos-mediated integration cassette), and PBac (PiggyBac) are the names of the respective insertions.

**Suppl. Table 3: Results of pairwise *z*-tests comparing $LC_{50}$s among resistant and susceptible *Drosophila suzukii* isofemale lines**

| Insecticide | Comparison | Estimate ± SE | *t* value | *p* value |
|---|---|---|---|---|
| Zeta-cypermethrin | S3 - S4 | 1.1646 ± 1.1081 | 1.5229 | 0.1278 |
| | S3 - S8 | 3.1909 ± 0.2962 | 7.3954 | 1.41e-13 |
| | S4 - S8 | 2.7398 ± 0.2760 | 6.3026 | 2.93e-10 |
| Spinosad | C3 - C4 | 1.2728 ± 0.1652 | 1.6512 | 0.0987 |
| | C3 - C5 | 6.7628 ± 0.9008 | 6.3974 | 1.58e-10 |
| | C4 - C5 | 5.3133 ± 0.7301 | 5.9078 | 3.47e-09 |

**Suppl. Table 23: Sequences for primers used in quantitative PCR**

| Primer Name | 5'- Sequence -3' |
|---|---|
| Ds*RpL32* (137) Forward | TGC GTC GCC GCT TCA AGG GAC |
| Ds*RpL32* (3287) Reverse | TGC GCT TCT TGG AGC TCA CGC C |
| Ds*Cyp6a8* (1049) Forward | TGA GGT GGA GGA TGT CCT AGA GC |
| Ds*Cyp6a8* (1215) Reverse | TCG GAT GGC CGG GAA CTT CG |
| Ds*Cyp4d14* (1589) Forward | TCC AGG AGA TTC GAG ATG TCC TTG |
| Ds*Cyp4d14* (1756) Reverse | TGC CGT CTA GCA CGG TGT CC |
| Ds*Cyp6w1* (1290) Forward | TCC GGC GAA CCG CTG TAA CCT C |
| Ds*Cyp6w1* (1479) Reverse | AAC CGG ACT AGT AGC AGC CCA C |
| Ds*TwdlG* (822) Forward | TCG CAC CCA AGC AAC CTA GCA AG |
| Ds*TwdlG* (999) Reverse | TGG TGG TCC AGA ACG CCA ATT AC |
| Ds*TwdlF* (842) Forward | AGC GCG CCC AGC AGG AGA |
| Ds*TwdlF* (1033) Reverse | AGC TCT GCT GCT GAA TGC CCT G |
| Ds*EcR* (2010) Forward | AGT CGC ACC TCC AGG TTA CA |
| Ds*EcR* (2174) Reverse | CGG TTG CGT ATT GTT TTG GGT |
| Dm*Cbp20* qPCR Forward | GTC TGA TTC GTG TGG ACT GG |
| Dm*Cbp20* qPCR Reverse | CAA CAG TTT GCC ATA ACC CC |
| Dm*Cyp4d8* (2238) Forward | TGG AGT AAT TCC GGC TGG CTC AG |
| Dm*Cyp4d8* (2431) Reverse | CTC CAG CTG AGC GAA CTT CTG AC |
| Dm*Cyp6d4* (1272) Forward | ACC TAC GAT TCC CTG AAC AAG ATG G |
| Dm*Cyp6d4* (1457) Reverse | TGC ATC ATG GTG AAT GCC GTA CAG |
| Dm*Cyp4d14* (1272) Forward | TCC AGG AGG TCA GAG ATG TTA TCG |
| Dm*Cyp4d14* (1457) Reverse | TGC CGT CGA GTA CGG TGT CC |
| Dm*Cyp6a20* (1211) Forward | ACA GAG ACC ATG AGG AAG CGT CC |
| Dm*Cyp6a20* (1355) Reverse | ACT CTG GAT CGT GAT GTA TGG CC |
| Dm*Cpr66D* (2674) Forward | AGG CCG AGG TCA TTC GCG AG |
| Dm*Cpr66D* (2853) Reverse | ACT GGT ACT GGT GGT ACT GTG GC |
